# The (Limited?) Utility of Brain Age as a Biomarker for Capturing Fluid Cognition in Older Individuals

**DOI:** 10.1101/2022.12.31.522374

**Authors:** Alina Tetereva, Narun Pat

**Affiliations:** Department of Psychology, University of Otago, New Zealand, 9016

**Keywords:** Brain Age, Fluid Cognition, brain MRI, machine learning, Aging, biomarker

## Abstract

Fluid cognition usually declines as people grow older. For decades, neuroscientists have been on a quest to search for a biomarker that can help capture fluid cognition. One well-known candidate is Brain Age, or a predicted value based on machine-learning models built to predict chronological age from brain MRI data. Here we aim to formally evaluate the utility of Brain Age as a biomarker for capturing fluid cognition among older individuals. Using 504 aging participants (36-100 years old) from the Human Connectome Project in Aging, we created 26 age-prediction models for Brain Age based on different combinations of MRI modalities. We first tested how much Brain Age from these age-prediction models added to what we had already known from a person’s chronological age in capturing fluid cognition. Based on the commonality analyses, we found a large degree of overlap between Brain Age and chronological age, so much so that, at best, Brain Age could uniquely add only around 1.6% in explaining variation in fluid cognition. Next, the age-prediction models that performed better at predicting chronological age did NOT necessarily create better Brain Age for capturing fluid cognition over and above chronological age. Instead, better-performing age-prediction models created Brain Age that overlapped larger with chronological age, up to around 29% out of 32%, in explaining fluid cognition, thus not improving the models’ utility to capture cognitive abilities. Lastly, we tested how much Brain Age missed the variation in the brain MRI that could explain fluid cognition. To capture this variation in the brain MRI that explained fluid cognition, we computed Brain Cognition, or a predicted value based on prediction models built to directly predict fluid cognition (as opposed to chronological age) from brain MRI data. We found that Brain Cognition captured up to an additional 11% of the total variation in fluid cognition that was missing from the model with only Brain Age and chronological age, leading to around a 1/3-time improvement of the total variation explained. Accordingly, we demonstrated the limited utility of Brain Age as a biomarker for fluid cognition and made some suggestions to ensure the utility of Brain Age in explaining fluid cognition and other phenotypes of interest.

## Introduction

Older adults often experience declines in several cognitive abilities such as memory, attention and processing speed, collectively known as fluid cognition (Salthouse, 2019; Weintraub et al., 2014). Having objective biomarkers to capture fluid cognition would give researchers and clinicians a tool to detect early cognitive impairments, monitor treatment/intervention efficacy and forecast cognitive prognosis (Frisoni et al., 2017). Over the past decade, Brain Age (Franke et al., 2010) has emerged as a potential biomarker to capture fluid cognition in older adults (Cole, 2020; Cole et al., 2018; Liem et al., 2017; Richard et al., 2018; Wrigglesworth et al., 2022; see review Boyle et al., 2021). Yet, to justify the use of Brain Age as an informative biomarker for fluid cognition, we still need to address at least the three impeding issues.

First, to what extent does having information on Brain Age improve its utility to capture fluid cognition in older adults beyond knowing a person’s chronological age? To compute Brain Age, researchers first build a prediction model that predicts the chronological age based on a person’s brain MRI data (Baecker et al., 2021). They then apply this prediction model to an unseen individual, not part of the model-building process. Brain Age is the predicted value of this model. Accordingly, by design, Brain Age is tightly close to chronological age. Because chronological age usually has a strong relationship with fluid cognition, to begin with, it is unclear how much Brain Age adds to what is already captured by chronological age.

Note researchers often subtract chronological age from Brain Age, creating an index known as Brain Age Gap (Franke & Gaser, 2019). A higher value of Brain Age Gap is thought to reflect accelerated/premature aging. Yet, given that Brain Age Gap is calculated based on both Brain Age and chronological age, Brain Age Gap still depends on chronological age (Butler et al., 2021). If, for instance, Brain Age was based on prediction models with poor performance and made a prediction that everyone was 50 years old, individual differences in Brain Age Gap would then depend solely on chronological age (i.e., 50 minus chronological age). Moreover, Brain Age is known to demonstrate the “regression towards the mean” phenomenon (Stigler, 1997). More specifically, because Brain Age is a predicted value of a regression model that predicts chronological age, Brain Age is usually shrunk towards the mean age of samples used for training the model (Butler et al., 2021; de Lange & Cole, 2020; Le et al., 2018). Accordingly, Brain Age predicts chronological age more accurately for individuals who are closer to the mean age while overestimating younger individuals’ chronological age and underestimating older individuals’ chronological age. There are many adjustments proposed to correct for the age dependency, but the outcomes tend to be similar to each other (Beheshti et al., 2019; de Lange & Cole, 2020; Liang et al., 2019; Smith et al., 2019). These adjustments can be applied to Brain Age and Brain Age Gap, creating Corrected Brain Age and Corrected Brain Age Gap, respectively. Corrected Brain Age Gap in particular is viewed as being able to control for age dependency (Butler et al., 2021). Here, we tested the utility of different Brain Age calculations in capturing fluid cognition, over and above chronological age.

Second, do better-performing age-prediction models correspond to the improvement in the utility to capture fluid cognition? Over the past decades, there has been a race to improve the performance of age-prediction models to be better at predicting chronological age, for instance, by combining different MRI/neuroimaging modalities features (Cole, 2020; Engemann et al., 2020; Liem et al., 2017) or by applying more sophisticated machine-learning algorithms (Baecker et al., 2021; Jonsson et al., 2019; Zhao & Zhao, 2021). However, the improvement in predicting chronological age may not necessarily make Brain Age Gap better at capturing fluid cognition (Jirsaraie, Gorelik, et al., 2023). If, for instance, the age-prediction model had the perfect performance, Brain Age Gap would be exactly zero and would have no utility in capturing fluid cognition beyond chronological age. Here we examined the performance of age-prediction models as a function of their utility in capturing fluid cognition.

Third and finally, certain variation in fluid cognition is related to brain MRI, but to what extent does Brain Age not capture this variation? To estimate the variation in fluid cognition that is related to brain MRI, we could build prediction models that directly predict fluid cognition (i.e., as opposed to chronological age) from brain MRI data. Previous studies found reasonable predictive performances of these cognition-prediction models, built from certain MRI modalities (Dubois et al., 2018; Pat, Wang, Anney, et al., 2022; Rasero et al., 2021; Sripada et al., 2020; Tetereva et al., 2022; for review, see Vieira et al., 2022). Analogous to Brain Age, we called the predicted values from these cognition-prediction models, Brain Cognition. The strength of an out-of-sample relationship between Brain Cognition and fluid cognition reflects variation in fluid cognition that is related to the brain MRI and, therefore, indicates the upper limit of Brain Age’s capability in capturing fluid cognition. This is, by design, the variation in fluid cognition explained by Brain Cognition should be higher or equal to that explained by Brain Age. Consequently, if we included Brain Cognition, Brain Age and chronological age in the same model to explain fluid cognition, we would be able to examine the unique effects of Brain Cognition that explain fluid cognition beyond Brain Age and chronological age. These unique effects of Brain Cognition, in turn, would indicate the amount of co-variation between brain MRI and fluid cognition that is missed by Brain Age.

Our study set out to test the utility of Brain Age as a biomarker for capturing variation in fluid cognition among aging individuals. Using aging participants (36-100 years old) from the Human Connectome Project in Aging (Bookheimer et al., 2019), we computed different Brain Age indices (including Brain Age, Brain Age Gap, Corrected Brain Age and Corrected Brain Age Gap) and Brain Cognition from prediction models based on different sets of MRI features. These MRI features covered task, resting-state and structural MRI, creating 26 prediction models in total. We, then, tested the biomarkers’ utility in explaining fluid cognition in unseen participants. To test this utility of Brain Age indices, we applied simple regression models with each Brain Age index as a sole regressor to explain fluid cognition. Next, to test the unique effects of Brain Age in explaining fluid cognition beyond chronological age, we applied multiple regression models with both each Brain Age index and chronological age as regressors to explain fluid cognition. To reveal how much chronological age and Brain Age indices had in common in explaining fluid cognition (i.e., common effects), we then applied the commonality analysis (Nimon et al., 2008) to these multiple regression models. Additionally, given that certain sets of MRI features led to prediction models that were better at predicting chronological age, we also examined if these better-performing age-prediction models improved the utility of Brain Age indices in explaining fluid cognition over and above lower-performing age-prediction models. Finally, we investigated the extent to which Brain Age indices missed the variation in fluid cognition that could be explained by the brain MRI. Here, we tested Brain Cognition’s unique effects in multiple regression models with a Brain Age index, chronological age and Brain Cognition as regressors to explain fluid cognition.

## Results

### Relationship between chronological age and fluid cognition

Figure 1a shows the negative relationship between chronological age and fluid cognition (*r*(502) = -.57, *p* < .001, R^2^ = .32). Older individuals tended to have a lower fluid cognition score.

**Figure 1.**
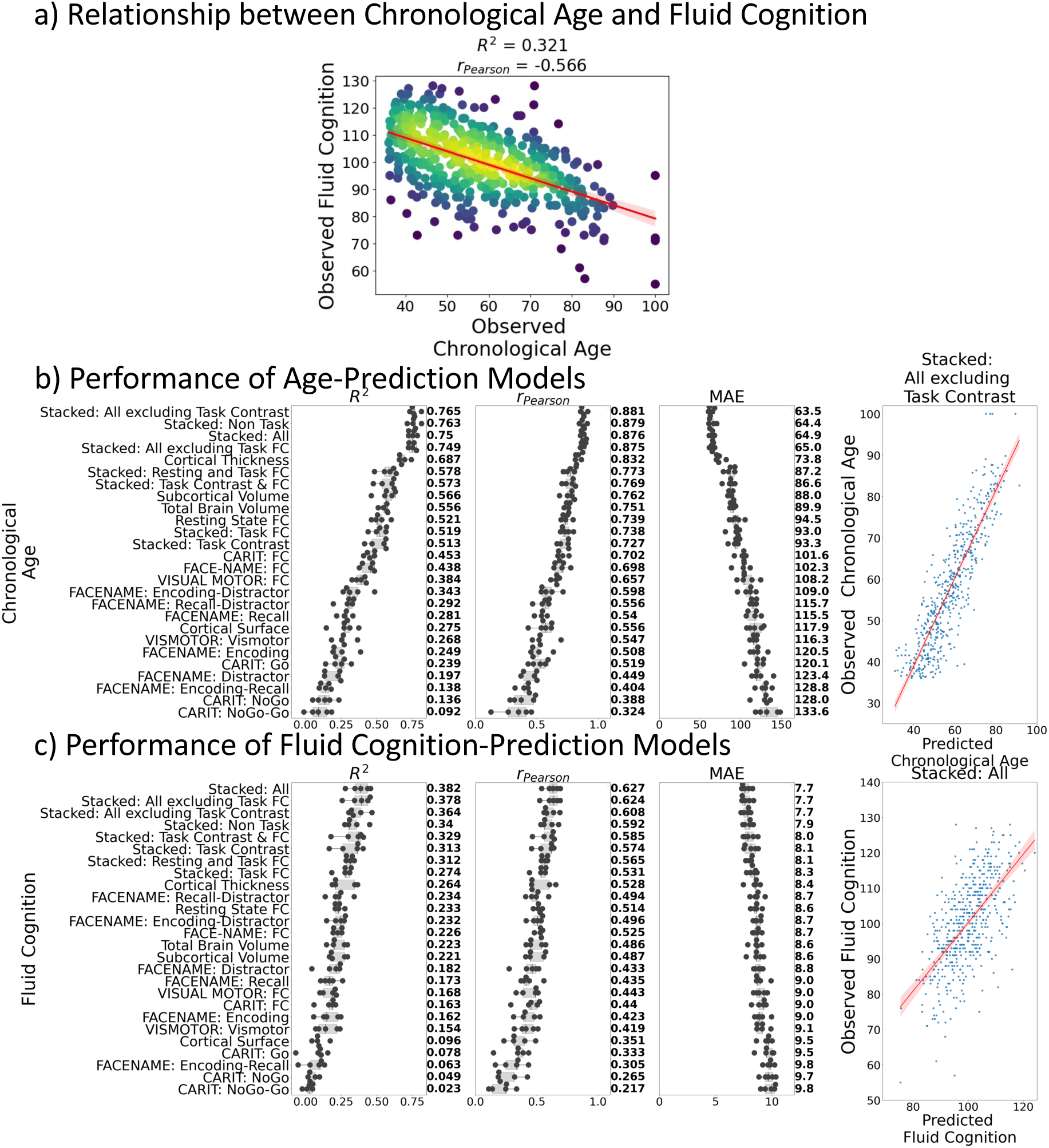
Relationship between chronological age and fluid cognition (a) and predictive performance of prediction models using Brain MRI from different sets of MRI features to predict chronological age (b) and fluid cognition (c). Each dot in (b) and (c) represents predictive performance at each of the five outer-fold test sets. The numbers to the right of the predictive performance plots indicate the mean of predictive performance across the five outer-fold test sets. Note we only provided the scatter plots between observed and predicted values in the outer-fold test sets from the best prediction models for each target (age in years and fluid cognition in points) in this figure. See Supplementary Figures 1 and 2 for the scatter plots from other prediction models.

### Predictive performance of prediction models for Brain Age and Brain Cognition

Figure 1b and 1c show the predictive performance of different sets of brain MRI features in predicting chronological age and fluid cognition, respectively. For age prediction, the top-four models that performed similarly were ‘stacked’ models that included multiple sets of brain MRI features: “Stacked: All excluding Task Contrast”, “Non Task”, “All excluding Task FC” and “All” (*R*^2^ > .76, *r* >.83, MAE < 65 months). For fluid cognition prediction, the top-performing model was “Stacked: All” (R^2^ = .393, *r* = .627, MAE = 7.7 points). The best set of features across age and fluid cognition prediction was cortical thickness. Across sets of MRI features, the age-prediction models tended to provide higher R^2^ and *r* than the fluid cognition-prediction models. Figure 2 shows the feature importance of prediction models based on each of the 18 sets of features. Figure 3 shows the feature importance of the eight stacked prediction models. Figure 4 shows the stability of feature importance across different outer-fold test sets.

**Figure 2.**
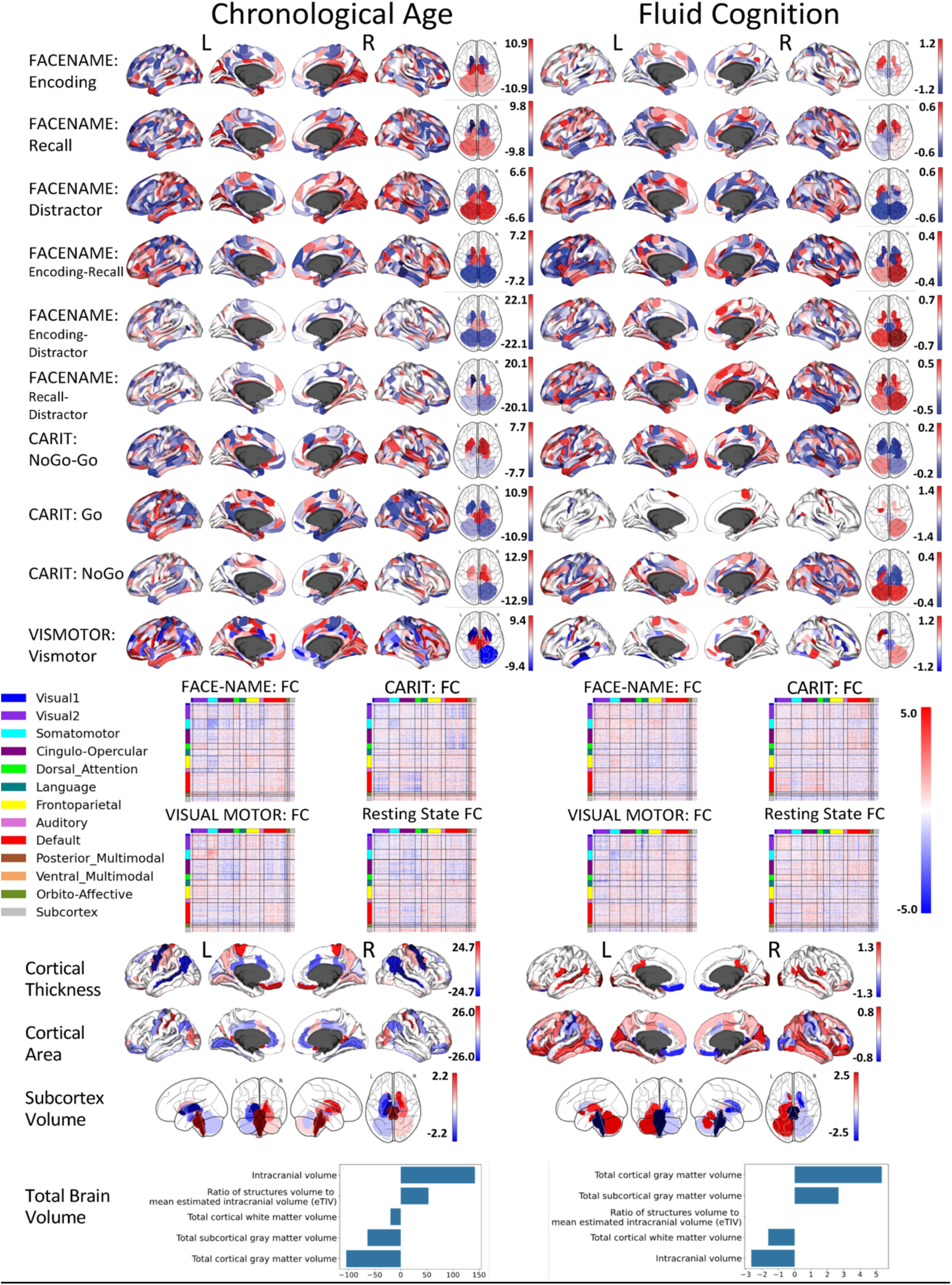
Feature importance (i.e., Elastic Net Coefficients) of prediction models based on each of the 18 sets of features.

**Figure 3.**
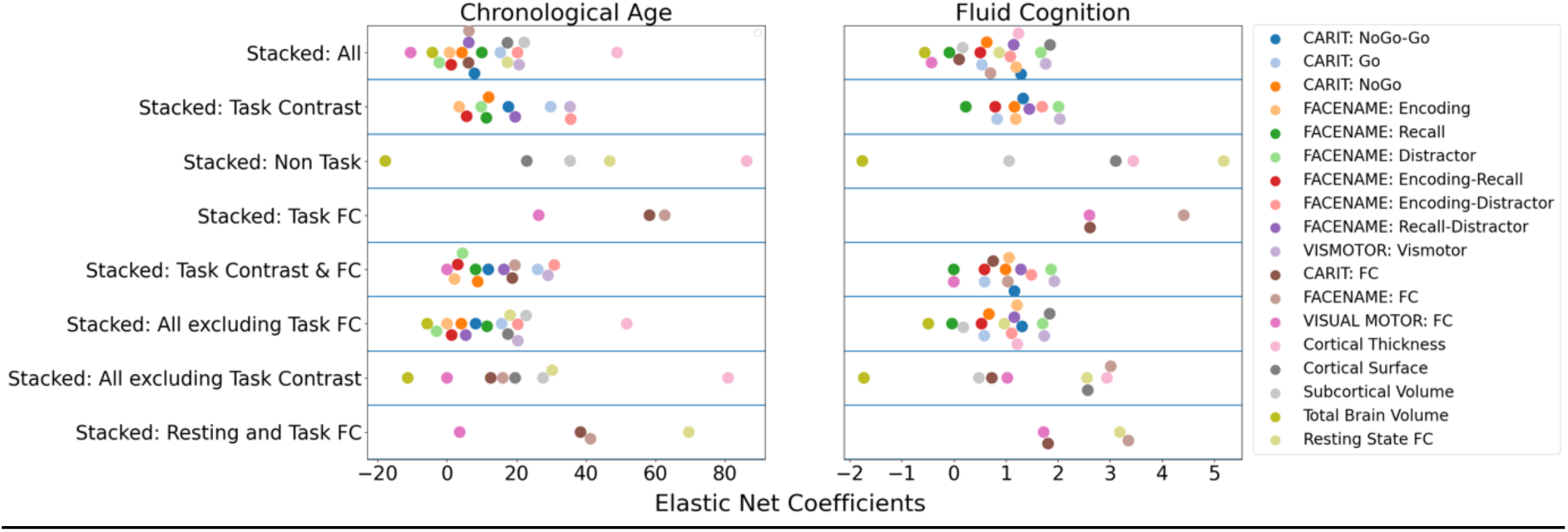
Feature importance (i.e., Elastic Net Coefficients) of the eight stacked prediction models.

**Figure 4.**
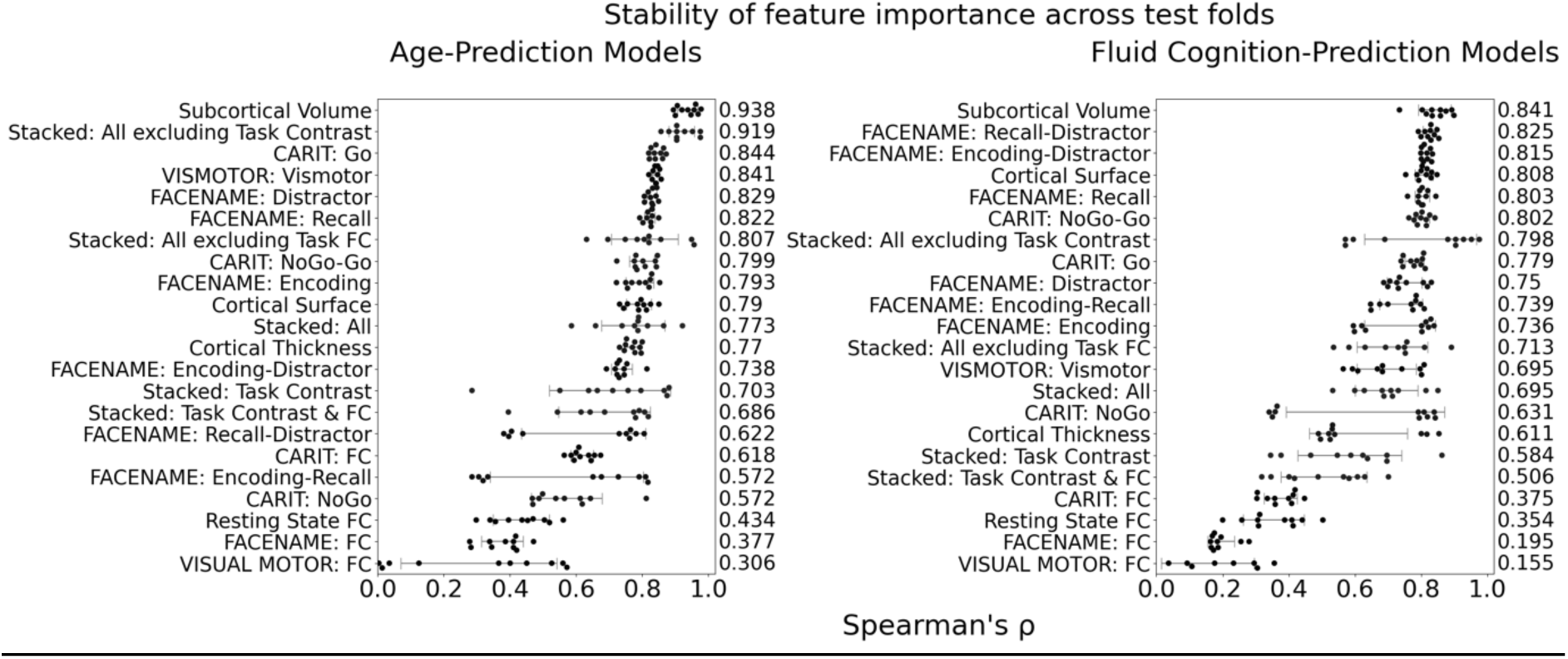
Stability of feature importance (i.e., Elastic Net Coefficients) of prediction models. Each dot represents rank stability (reflected by Spearman’s ρ) in the feature importance between two prediction models of the same features, used in two different outer-fold test sets. Given that there were five outer-fold test sets, there were 10 Spearman’s ρs for each prediction model. The numbers to the right of the plots indicate the mean of Spearman’s ρ for each prediction model.

### Simple regression: Using each Brain Age index to explain fluid cognition

Figure 5a shows variation in fluid cognition explained by Brain Age Indices when having each Brain Age index as the sole regressor in simple regression models. Brain Age and Corrected Brain Age created from higher-performing age-prediction models explained a higher amount of variation in fluid cognition. However, Brain Age Gap created from the *lower*-performing age-prediction models explained a higher amount of variation in fluid cognition. For instance, the top performing age-prediction model, “Stacked: All excluding Task Contrast”, generated Brain Age and Corrected Brain Age that explained the highest amount of variation in fluid cognition, but, at the same time, produced Brain Age Gap that explained the least amount of variation in fluid cognition.

**Figure 5.**
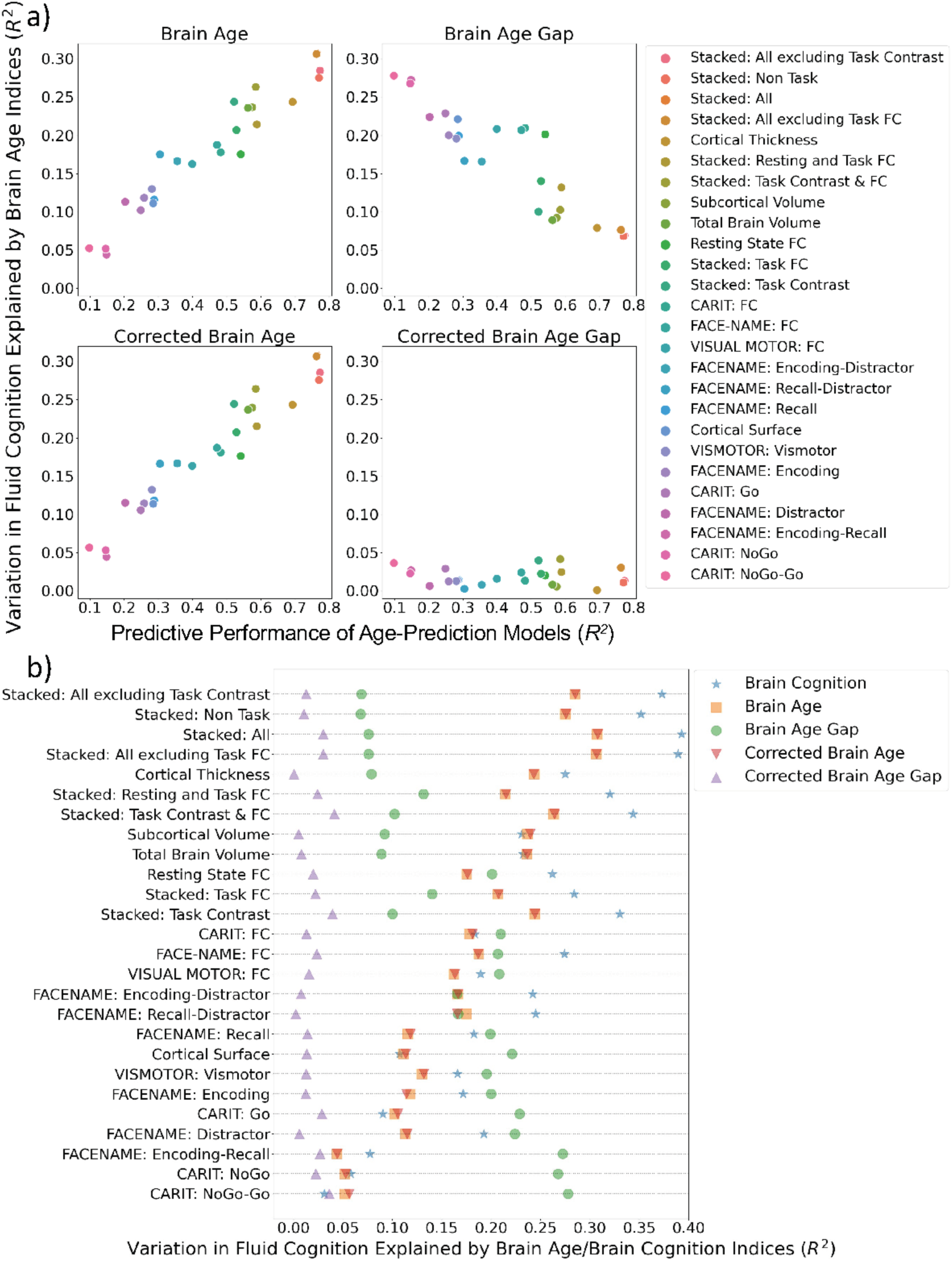
Simple regression: using each Brain Age index or Brain Cognition to explain fluid cognition. 5a shows variation in fluid cognition explained by each Brain Age index as a function of the predictive performance of age-prediction models. 5b plots variation in fluid cognition explained by Brain Age indices and Brain Cognition.

On the contrary, an amount of variation in fluid cognition explained by Corrected Brain Age Gap was relatively small (maximum at *R*^2^=.041) across age-prediction models and did not relate to the predictive performance of the age-prediction models. Figure 5b shows variation in fluid cognition explained by Brain Cognition, as compared to Brain Age indices. Brain Cognition appeared to explain a higher amount of variation in fluid cognition than any Brain Age indices, especially for top-performing age/cognition-prediction models (e.g., Stacked: All).

### Multiple regression: Using chronological age and each Brain Age index to explain fluid cognition

Figure 6 shows the commonality analysis of multiple regression models, having both chronological age and each Brain Age index as the regressors for fluid cognition. We found *R*^2^ for these models at *M* = .326 (SD = 005). The unique effects of Brain Age indices were all relatively small (maximum at Δ*R*^2^Brain Age index = .0161, with statistically significant at *p-value* < .05 in 10 out of 26 models) across the four Brain Age indices and across different age-prediction models.

**Figure 6.**
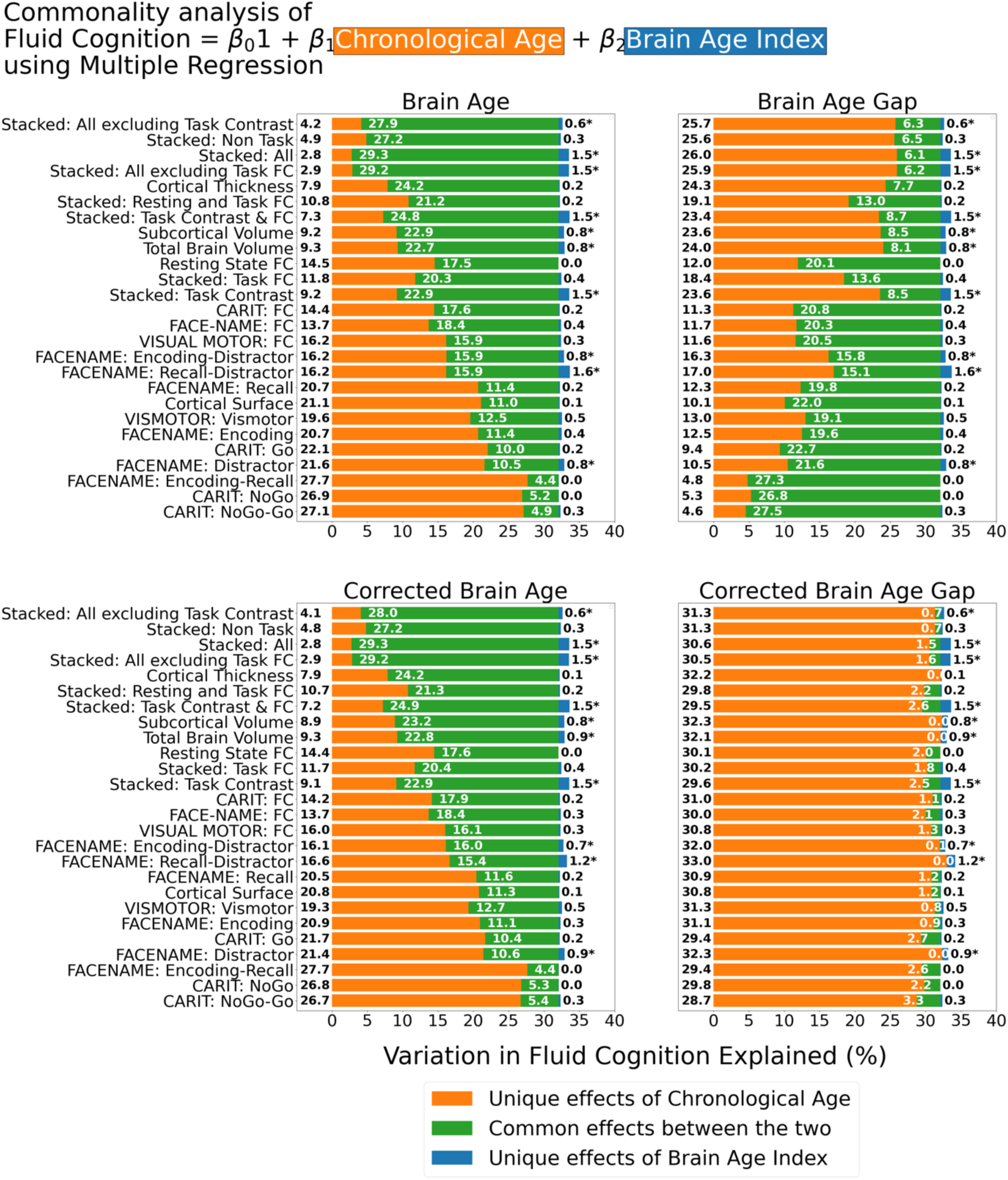
Commonality analysis of multiple regressions, having both chronological age and each Brain Age index as the regressors for capturing fluid cognition. The numbers to the left of the figure represent the unique effects of chronological age in %, the numbers in the middle of the figure represent the common effects between chronological age and Brain Age index in %, and the numbers to the right of the figure represent the unique effects of Brain Age Index in %. * represents the statistical significance of the unique effects of Brain Age Index at p < .05.

However, it is clear that different Brain Age indices led to different levels of the unique effects of chronological age and the common effects between chronological age and Brain Age indices. For the top-performing age-prediction models (e.g., Stacked: All excluding Task Contrast), the unique effects of chronological age were low for Brain Age and Corrected Brain Age, but high for Brain Age Gap. On the contrary, the lower-performing age-prediction models provided high common effects for Brain Age and Corrected Brain Age, but low for Brain Age Gap. Nonetheless, for Corrected Brain Age Gap, the unique effects of chronological age were much higher than the common effects across all age-prediction models.

### Multiple regression: Using chronological age, each Brain Age index and Brain Cognition to explain fluid cognition

Figure 7 shows the commonality analysis of multiple regression models, having chronological age, each Brain Age index and Brain Cognition as the regressors for fluid cognition. We found *R*^2^ for these models at *M*=.385 (SD=.042). As before, the unique effects of Brain Age indices were all relatively small across the four Brain Age indices and across different prediction models. On the contrary, the unique effects of Brain Cognition appeared much larger (maximum at Δ*R*^2^_cognition_ = .1183, statistically significant *p-value* at .05 in 24 out of 26 models).

**Figure 7.**
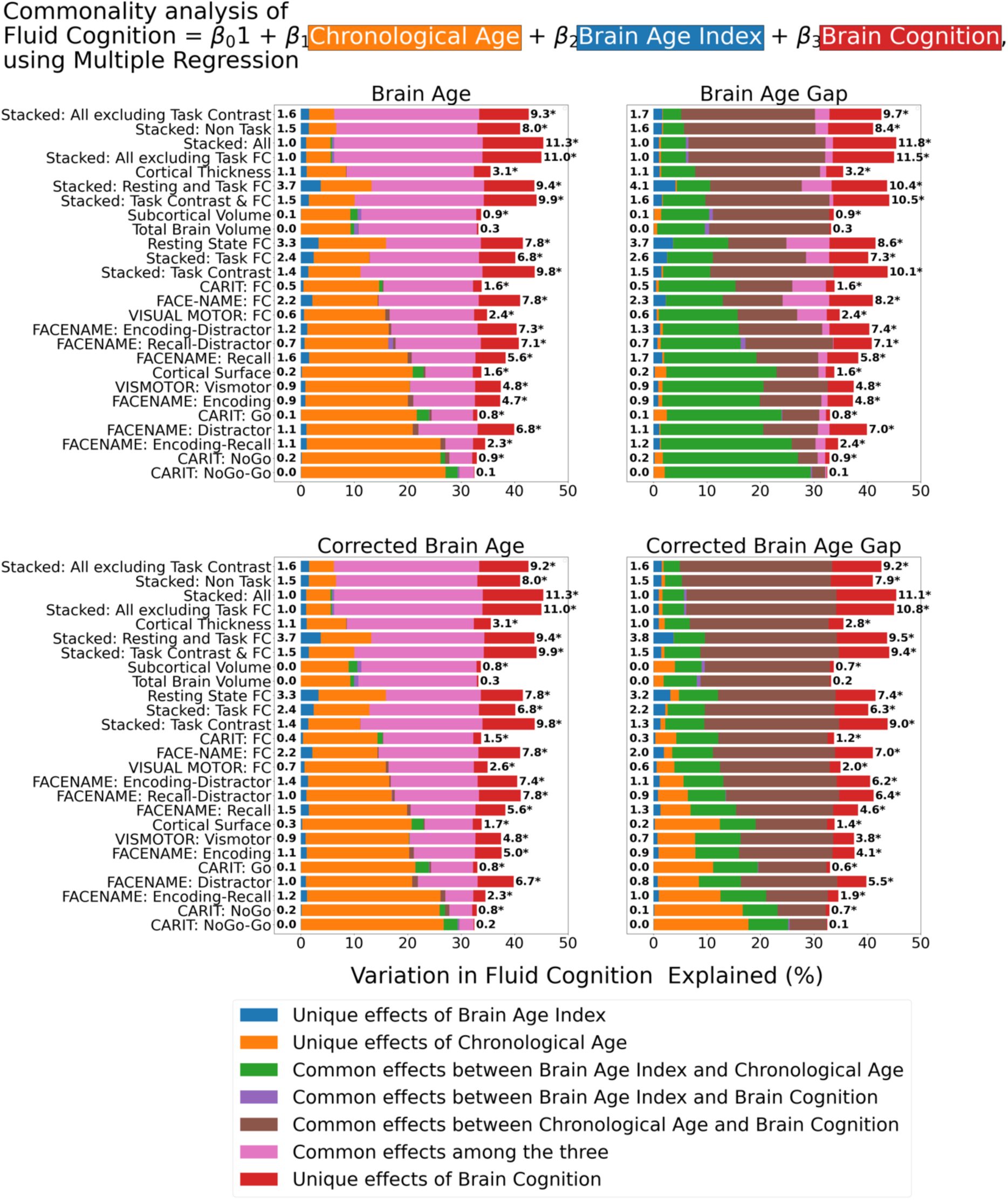
Commonality analysis of multiple regressions, having chronological age and each Brain Age index and Brain Cognition as the regressors for capturing fluid cognition. The numbers to the left of the figures represent the unique effects of Brain Age Index in %, and the numbers to the right of the figures represent the unique effects of Brain Cognition in %. * represents the statistical significance of the unique effects of Brain Cognition at p < .05.

For top-performing age/cognition-prediction models (e.g., Stacked All), the largest proportion of fluid cognition was attributed to a) the common effects among the three for Brain Age and Corrected Brain Age and b) the common effects between chronological age and Brain Cognition for Brain Age Gap and Corrected Brain Age Gap.

## Discussion

To demonstrate the utility of Brain Age as a biomarker for fluid cognition, we investigated three essential issues. First, how much does Brain Age add to what is already captured by chronological age? The short answer is very little. Second, do better-performing age-prediction models improve the utility of Brain Age to capture fluid cognition above and beyond chronological age? The answer is also no. Third, how much does Brain Age miss the variation in the brain MRI that could explain fluid cognition? Brain Age and chronological age by themselves captured around 32% of the total variation in fluid cognition. But, around an additional 11% of the variation in fluid cognition could have been captured if we used the prediction models that directly predicted fluid cognition from brain MRI.

First, Brain Age itself did not add much more information to help us capture fluid cognition than what we had already known from a person’s chronological age. This can clearly be seen from the small unique effects of Brain Age indices in the multiple regression models having Brain Age and chronological age as the regressors. While the unique effects of some Brain Age indices from certain age-prediction models were statistically significant, there were all relatively small. Without Brain Age indices, chronological age by itself already explained around 32% of the variation in fluid cognition. Including Brain Age indices only added around 1.6% at best. We believe the small unique effects of Brain Age were inevitable because, by design, Brain Age is tightly close to chronological age. Therefore, chronological age and Brain Age captured mostly a similar variation in fluid cognition.

Investigating the simple regression models and the commonality analysis between each Brain Age index and chronological age provided additional insights. In the simple regression models, higher-performing age-prediction models, such as stacked models, created Brain Age and Corrected Brain Age that captured a higher amount of variation in fluid cognition. Because both Brain Age and Corrected Brain Age from higher-performing age-prediction models were closer to the real chronological age of participants, their ability to capture fluid cognition mirrored the ability of chronological age. The commonality analysis confirmed this by showing higher common effects between (Corrected) Brain Age and chronological age from higher-performing age-prediction models. In contrast, lower-performing (as opposed to higher-performing) age-prediction models, such as CARIT NoGo-Go, created Brain Age Gap that explained a higher amount of variation in fluid cognition. Brain Age Gap was a result of subtracting a real chronological age from Brain Age. And when Brain Age was a poor indicator of the real chronological age, the utility of Brain Age Gap is driven more by the real chronological age (Butler et al., 2021). The commonality analysis confirmed this by showing higher common effects, therefore more similarity in variance, between Brain Age Gap and chronological age from lower-performing, than higher-performing, age-prediction models.

Corrected Brain Age Gap, on the other hand, showed weak effects on the simple regression models across all age-prediction models (max at around 4.1% of variation explained). Corrected Brain Age Gap was the only index among the four that appeared to deconfound the influences of chronological age on the relationship between brain aging and fluid cognition (Butler et al., 2021). In our study, this can be seen in the small common effects between Corrected Brain Age Gap and chronological age in the multiple regression models with chronological age and each Brain Age index as regressors. Note while these common effects between Corrected Brain Age Gap and chronological age were small, most were not zero (max at around 3.3% of variation explained). This means that the correction done to deconfound the influences of chronological age on Corrected Brain Age Gap (de Lange & Cole, 2020) may not be perfect. Perhaps this is because the estimation of the influences of chronological age was done in the training set, which might not fully be applicable to the test sets. Still, weak effects of Corrected Brain Age Gap in the simple regression indicate that, after controlling for the influences of chronological age, this Brain Age index could only account for a small amount of variation in fluid cognition. In other words, the weak effects of Corrected Brain Age Gap shown by the simple regression are consistent with the small unique effects across the four Brain Age indices shown by the multiple regression models having a Brain Age index and chronological age as regressors.

The small effects of the Corrected Brain Age Gap in explaining fluid cognition of aging individuals found here are consistent with studies in older adults (Cole, 2020) and younger populations (Butler et al., 2021; Jirsaraie, Kaufmann, et al., 2023). Cole (2020) studied the utility of Brain Age on cognitive functioning of large samples (n>17,000) of older adults, aged 45-80 years, from the UK Biobank (Sudlow et al., 2015). He constructed age-prediction models using LASSO, a similar penalised regression to ours and applied the same age-dependency adjustment to ours. Cole (2020) then conducted a multiple regression explaining cognitive functioning from Corrected Brain Age Gap while controlling for chronological age and other potential confounds. He found Corrected Brain Age Gap to be significantly related to performance in four out of six cognitive measures, and among those significant relationships, the effect sizes were small with a maximum of partial eta-squared at .0059. Similarly, Jirsaraie and colleagues (2023) studied the utility of Brain Age on cognitive functioning of youths aged 8-22 years old from the Human Connectome Project in Development (Somerville et al., 2018) and Preschool Depression Study (Luby, 2010). They built age-prediction models using gradient tree boosting (GTB) and deep-learning brain network (DBN) and adjusted the age dependency of Brain Age Gap using Smith and colleagues’ (2019) method. Using multiple regressions, Jirsaraie and colleagues (2023) found weak effects of the adjusted Brain Age Gap on cognitive functioning across five cognitive tasks, five age-prediction models and the two datasets (mean of standardised regression coefficient = -0.09, see their Table S7). Next, Butler and colleagues (2021) studied the utility of Brain Age on cognitive functioning of another group of youths aged 8-22 years old from the Philadelphia Neurodevelopmental Cohort (PNC) (Satterthwaite et al., 2016). Here they used Elastic Net to build age-prediction models and applied another age-dependency adjustment method, proposed by Beheshti and colleagues (2019). Similar to the aforementioned results, Butler and colleagues (2021) found a weak, statistically non-significant correlation between the adjusted Brain Age Gap and cognitive functioning at *r*=-.01, *p*=.71. Accordingly, the utility of Brain Age in explaining cognitive functioning beyond chronological age appears to be weak across age groups, different predictive modelling algorithms and age-dependency adjustments.

Second, the predictive performance of age-prediction models did not correspond to the utility of Brain Age in capturing fluid cognition over and above chronological age. For instance, while the best-performing age-prediction model was “Stacked: All excluding Task Contrast” (*R*^2^=.775), the unique effects of Brain Age indices from this model in the two-regressor multiple regressions (i.e., with a Brain Age index and chronological age as regressor) were weak (Δ*R*^2^_Brain Age index_ ≤.0048) and not statistically significant. The highest unique effects of Brain Age indices in the two-regressor multiple regression models were from the FACENAME: Distractor model (Δ*R*^2^_Brain Age index_ ≤.0135, *p* < .05) that had a poorer performance in predicting chronological age (*R*^2^=.204). Accordingly, a race to improve the performance of age-prediction models (Baecker et al., 2021) does not necessarily enhance the utility of Brain Age indices as a biomarker for fluid cognition. This discrepancy between the predictive performance of age-prediction models and the utility of Brain Age indices as a biomarker is consistent with recent findings (for review, see Jirsaraie, Gorelik, et al., 2023), both in the context of cognitive functioning (Jirsaraie, Kaufmann, et al., 2023) and neurological/psychological disorders (Bashyam et al., 2020; Rokicki et al., 2021). For instance, combining different MRI modalities into the prediction models, similar to our stacked models, often leads to the highest performance of age-prediction models, but does not likely explain the highest variance across different phenotypes, including cognitive functioning and beyond (Jirsaraie, Gorelik, et al., 2023).

Third, by introducing Brain Cognition, we showed the extent to which Brain Age indices were not able to capture the variation in fluid cognition that is related to brain MRI. More specifically, using Brain Cognition allowed us to gauge the variation in fluid cognition that is related to the brain MRI, and thereby, to estimate the upper limit of what Brain Age can do. Moreover, by examining what was captured by Brain Cognition, over and above Brain Age and chronological age via the unique effects of Brain Cognition, we were able to quantify the amount of co-variation between brain MRI and fluid cognition that was missed by Brain Age.

From our results, Brain Cognition, especially from certain cognition-prediction models such as the stacked models, has relatively good predictive performance, consistent with previous studies (Dubois et al., 2018; Pat, Wang, Anney, et al., 2022; Rasero et al., 2021; Sripada et al., 2020; Tetereva et al., 2022; for review, see Vieira et al., 2022). We then examined Brain Cognition using commonality analyses (Nimon et al., 2008) in multiple regression models having a Brain Age index, chronological age and Brain Cognition as regressors to explain fluid cognition. Similar to Brain Age indices, Brain Cognition exhibited large common effects with chronological age. But more importantly, unlike Brain Age indices, Brain Cognition showed large unique effects, up to around 11%. As explained above, the unique effects of Brain Cognition indicated the amount of co-variation between brain MRI and fluid cognition that was missed by a Brain Age index and chronological age. This missing amount was relatively high, considering that Brain Age and chronological age together explained around 32% of the total variation in fluid cognition. Accordingly, if a Brain Age index was used as a biomarker along with chronological age, we would have missed an opportunity to improve the performance of the model by around one-third of the variation explained.

There are several potential limitations of this study. First, we conducted an investigation relying only on one dataset, the Human Connectome Project in Aging (HCP-A) (Bookheimer et al., 2019). While HCP-A used state-of-the-art MRI methodologies, covered a wide age range from 36 to 100 years old and used several task-fMRI from different tasks that are harder to find in other bigger databases (e.g., UK Biobank from Sudlow et al., 2015), several characteristics of HCP-A might limit the generalisability of our findings. For instance, the tasks used in task-based fMRI in HCP-A are not used widely in clinical settings (Horien et al., 2020). This might make it challenging to translate the approaches used here. Similarly, HCP-A also excluded participants with neurological conditions, possibly making their participants not representative of the general population. Next, while HCP-A’s sample size is not small (n=725 and 504 people, before and after exclusion, respectively), other datasets provide a much larger sample size (Horien et al., 2020). Similarly, HCP-A does not include younger populations. But as mentioned above, a study with a larger sample in older adults (Cole, 2020) and studies in younger populations (8-22 years old) (Butler et al., 2021; Jirsaraie, Kaufmann, et al., 2023) also found small effects of the adjusted Brain Age Gap in explaining cognitive functioning. And the disagreement between the predictive performance of age-prediction models and the utility of Brain Age found here is largely in line with the findings across different phenotypes seen in a recent systematic review (Jirsaraie, Gorelik, et al., 2023).

There is a notable difference between studies investigating the utility of Brain Age in explaining cognitive functioning, including ours and others (e.g., Butler et al., 2021; Cole, 2020, 2020; Jirsaraie, Kaufmann, et al., 2023) and those explaining neurological/psychological disorders (e.g., Bashyam et al., 2020; Rokicki et al., 2021). We consider the former as a normative type of study and the latter as a case-control type of study (Insel et al., 2010; Marquand et al., 2016). Those case-control Brain Age studies focusing on neurological/psychological disorders often build age-prediction models from MRI data of largely healthy participants (e.g., controls in a case-control design or large samples in a population-based design), apply the built age-prediction models to participants without vs. with neurological/psychological disorders and compare Brain Age indices between the two groups. On the one hand, this means that case-control studies treat Brain Age as a method to detect anomalies in the neurological/psychological group (Hahn et al., 2021). On the other hand, this also means that case-control studies have to ignore under-fitted models when applied prediction models built from largely healthy participants to participants with neurological/psychological disorders (i.e., Brain Age may predict chronological age well for the controls, but not for those with a disorder). On the contrary, our study and other normative studies focusing on cognitive functioning often build age-prediction models from MRI data of largely healthy participants and apply the built age-prediction models to participants who are also largely healthy. Accordingly, the age-prediction models for explaining cognitive functioning in normative studies, while not allowing us to detect group-level anomalies, do not suffer from being under-fitted. This unfortunately might limit the generalisability of our study into just the normative type of study. Future work is still needed to test the utility of brain age in the case-control case.

What does it mean then for researchers/clinicians who would like to use Brain Age as a biomarker? First, they have to be aware of the overlap in variation between Brain Age and chronological age and should focus on the contribution of Brain Age over and above chronological age. Using Brain Age Gap will not fix this. Butler and colleagues (2021) recently highlighted this point, “These results indicate that the association between cognition and the BAG [Brain Age Gap] are driven by the association between age and cognitive performance. As such, it is critical that readers of past literature note whether or not age was controlled for when testing for effects on the BAG, as this has not always been common practice (*p*. 4097).” Similar to previous recommendations (Butler et al., 2021; Le et al., 2018), we suggest future work should account for the relationship between Brain Age and chronological age, either using Corrected Brain Age Gap (or other similar adjustments) or, better, examining unique effects of Brain Age indices after controlling for chronological age through commonality analyses. Note we prefer using the commonality analysis as it can decompose variance of the phenotype of interest into unique and common effects, allowing us to understand the shared variance between chronological age and Brain Age indices (Ray-Mukherjee et al., 2014). In our case, Brain Age indices had the same unique effects regardless of the level of common effects they had with chronological age (e.g., Brain Age vs. Corrected Brain Age Gap from stacked models). In the case of fluid cognition, the unique effects might be too small to be clinically meaningful as shown here and previously (Butler et al., 2021; Cole, 2020; Jirsaraie, Kaufmann, et al., 2023).

Next, researchers should not select age-prediction models based solely on age-prediction performance. Instead, researchers could select age-prediction models that explained phenotypes of interest the best. Here we selected age-prediction models based on a set of features (i.e., modalities) of brain MRI. This strategy was found effective not only for fluid cognition as we demonstrated here, but also for neurological and psychological disorders as shown elsewhere (Jirsaraie, Gorelik, et al., 2023; Rokicki et al., 2021). Rokicki and colleagues (2021), for instance, found that, while integrating across MRI modalities led to age-prediction models with the highest age-prediction performance, using only T1 structural MRI gave age-prediction models that were better at classifying Alzheimer’s disease. Similarly, using only cerebral blood flow gave age-prediction models that were better at classifying mild/subjective cognitive impairment, schizophrenia and bipolar disorder.

As opposed to selecting age-prediction models based on a set of features, researchers could also select age-prediction models based on modelling methods. For instance, Jirsaraie and colleagues (2023) compared gradient tree boosting (GTB) and deep-learning brain network (DBN) algorithms in building age-prediction models. They found GTB to have higher age-prediction performance but DBN to have better utility in explaining cognitive functioning. In this case, an algorithm with better utility (e.g., DBN) should be used for explaining a phenotype of interest. Bashyam and colleagues (2020) made a similar observation, though for a contradictory conclusion, see Hahn and colleagues’ (2021). Bashyam and colleagues built different DBN-based age-prediction models, varying in age-prediction performance. The DBN models with a higher number of epochs corresponded to higher age-prediction performance. However, DBN-based age-prediction models with a moderate (as opposed to higher or lower) number of epochs were better at classifying Alzheimer’s disease, mild cognitive impairment and schizophrenia. In this case, a model from the same algorithm with better utility (e.g., those DBN with a moderate epoch number) should be used for explaining a phenotype of interest. In any case, this calls for a change in research practice, as recently pointed out by Jirasarie and colleagues (2023, p7), “Despite mounting evidence, there is a persisting assumption across several studies that the most accurate brain age models will have the most potential for detecting differences in a given phenotype of interest”. Future neuroimaging research should aim to build age-prediction models that are not necessarily good at predicting age, but at capturing phenotypes of interest.

Finally, researchers should test how much Brain Age miss the variation in the brain MRI that could explain fluid cognition or other phenotypes of interest. As demonstrated here, one straightforward method is to build a prediction model using a phenotype of interest as the target (e.g., fluid cognition) and incorporate the predicted value of this model (e.g., Brain Cognition), along with Brain Age and chronological age, into a multiple regression for commonality analyses. The unique effect of this predicted value will inform the missing variation in the brain MRI from Brain Age. If this unique effect is large, then researchers might need to reconsider whether using Brain Age is appropriate for a particular phenotype of interest.

Altogether, we examined the utility of Brain Age as a biomarker for fluid cognition. Here are the three conclusions. First, Brain Age failed to add substantially more information over and above chronological age. Second, a higher ability to predict chronological age did not correspond to a higher utility to capture fluid cognition. Third, Brain Age missed up to around one-third of the variation in fluid cognition that could have been explained by brain MRI. Yet, given our focus on fluid cognition, future empirical research is needed to test the utility of Brain Age on other phenotypes, especially when Brain Age is used for anomaly detection in case-control studies (e.g., Bashyam et al., 2020; Rokicki et al., 2021). We hope that future studies may consider applying our approach (i.e., using the commonality analysis that includes predicted values from a model that directly predicts the phenotype of interest) to test the utility of Brain Age as a biomarker for other phenotypes.

## Methods and Materials

### Dataset

We used the Human Connectome Project in Aging (HCP-A) (Bookheimer et al., 2019) Release 2.0 (24-February-2021). HCP-A’s ‘typical-aging’ participants (36-100 years old) may have prevalent health conditions (e.g., hypertension and different forms of vascular risks) but did not have identified pathological causes of cognitive decline (e.g., stroke and clinical dementia). In this Release, HCP-A provided data from 725 participants. HCP-A offered quality control flags, and here, we removed participants with the flag ‘A‘ anatomical anomalies or ‘B’ segmentation and surface (n= 117). Following further removal of participants with missing values in any of MRI modalities (n=15) or cognitive measurements (n= 111), we ultimately included 504 individuals (293 females, M= 57.83 (SD=14.25) years old) in our analyses. Note there were four individuals who were over 90 years old. HCP-A coded the age of these 90+ individuals as 100 years old to reduce the leakage of their personal health information. (See https://groups.google.com/a/humanconnectome.org/g/hcp-users/c/esZTVCRuxwE/m/xx4PLYMlCQAJ). For ethical procedures including informed consent, please see Bookheimer and colleagues’ (2019).

### Sets of brain MRI features

HCP-A provides details of parameters for brain MRI elsewhere (Bookheimer et al., 2019; Harms et al., 2018). Here we used MRI data that were pre-processed by the HCP-A with recommended methods, including the MSMALL alignment (Glasser et al., 2016; Robinson et al., 2018) and ICA-FIX (Glasser et al., 2016) for functional MRI. We used multiple brain MRI modalities, covering task functional MRI (task fMRI), resting-state functional MRI (rsfMRI) and structural MRI (sMRI), and organised them into 19 sets of features.

#### Sets of Features 1-10: Task fMRI contrast (Task Contrast)

Task contrasts reflect fMRI activation relevant to events in each task. Bookheimer and colleagues (2019) provided detailed information about the fMRI in HCP-A. Here we focused on the pre-processed task fMRI Connectivity Informatics Technology Initiative (CIFTI) files with a suffix, “_PA_Atlas_MSMAll_hp0_clean.dtseries.nii.” These CIFTI files encompassed both the cortical mesh surface and subcortical volume (Glasser et al., 2013). Collected using the posterior-to-anterior (PA) phase, these files were aligned using MSMALL (Glasser et al., 2016; Robinson et al., 2018), linear detrended (see https://groups.google.com/a/humanconnectome.org/g/hcp-users/c/ZLJc092h980/m/GiihzQAUAwAJ) and cleaned from potential artifacts using ICA-FIX (Glasser et al., 2016).

To extract Task Contrasts, we regressed the fMRI time series on the convolved task events using a double-gamma canonical hemodynamic response function via FMRIB Software Library (FSL)’s FMRI Expert Analysis Tool (FEAT) (Woolrich et al., 2001). We kept FSL’s default high pass cutoff at 200s (i.e., .005 Hz). We then parcellated the contrast ‘cope’ files, using the Glasser atlas (Gordon et al., 2016) for cortical surface regions and the Freesurfer’s automatic segmentation (aseg) (Fischl et al., 2002) for subcortical regions. This resulted in 379 regions, whose number was, in turn, the number of features for each Task Contrast set of features.

HCP-A collected fMRI data from three tasks: Face Name (Sperling et al., 2001), Conditioned Approach Response Inhibition Task (CARIT) (Somerville et al., 2018) and VISual MOTOR (VISMOTOR) (Ances et al., 2009).

First, the Face Name task (Sperling et al., 2001) taps into episodic memory. The task had three blocks. In the encoding block [Encoding], participants were asked to memorise the names of faces shown. These faces were then shown again in the recall block [Recall] when the participants were asked if they could remember the names of the previously shown faces. There was also the distractor block [Distractor] occurring between the encoding and recall blocks. Here participants were distracted by a Go/NoGo task. We computed six contrasts for this Face Name task: [Encode], [Recall], [Distractor], [Encode vs. Distractor], [Recall vs. Distractor] and [Encode vs. Recall].

Second, the CARIT task (Somerville et al., 2018) was adapted from the classic Go/NoGo task and taps into inhibitory control. Participants were asked to press a button to all [Go] but not to two [NoGo] shapes. We computed three contrasts for the CARIT task: [NoGo], [Go] and [NoGo vs. Go].

Third, the VISMOTOR task (Ances et al., 2009) was designed to test simple activation of the motor and visual cortices. Participants saw a checkerboard with a red square either on the left or right. They needed to press a corresponding key to indicate the location of the red square. We computed just one contrast for the VISMOTOR task: [Vismotor], which indicates the presence of the checkerboard vs. baseline.

#### Sets of Features 11-13: Task fMRI functional connectivity (Task FC)

Task FC reflects functional connectivity (FC) among the brain regions during each task, which is considered an important source of individual differences (Elliott et al., 2019; Fair et al., 2007; Gratton et al., 2018). We used the same CIFTI file “_PA_Atlas_MSMAll_hp0_clean.dtseries.nii.” as the task contrasts. Unlike Task Contrasts, here we treated the double-gamma, convolved task events as regressors of no interest and focused on the residuals of the regression from each task (Fair et al., 2007). We computed these regressors on FSL, and regressed them in nilearn (Abraham et al., 2014). Following previous work on task FC (Elliott et al., 2019), we applied a highpass at .008 Hz. For parcellation, we used the same atlases as Task Contrast (Fischl et al., 2002; Glasser et al., 2016). We computed Pearson’s correlations of each pair of 379 regions, resulting in a table of 71,631 non-overlapping FC indices for each task. We then applied r-to-z transformation and principal component analysis (PCA) of 75 components (Rasero et al., 2021; Sripada et al., 2019, 2020). Note to avoid data leakage, we conducted the PCA on each training set and applied its definition to the corresponding test set. Accordingly, there were three sets of 75 features for Task FC, one for each task.

#### Set of Features 14: Resting-state functional MRI functional connectivity (Rest FC)

Similar to Task FC, Rest FC reflects functional connectivity (FC) among the brain regions, except that Rest FC occurred during the resting (as opposed to task-performing) period. HCP-A collected Rest FC from four 6.42-min (488 frames) runs across two days, leading to 26-min long data (Harms et al., 2018). On each day, the study scanned two runs of Rest FC, starting with anterior-to-posterior (AP) and then with posterior-to-anterior (PA) phase encoding polarity. We used the “rfMRI_REST_Atlas_MSMAll_hp0_clean.dscalar.nii” file that was pre-processed and concatenated across the four runs. We applied the same computations (i.e., highpass filter, parcellation, Pearson’s correlations, r-to-z transformation and PCA) with the Task FC.

#### Sets of Features 15-18: Structural MRI (sMRI)

sMRI reflects individual differences in brain anatomy. The HCP-A used an established pre-processing pipeline for sMRI (Glasser et al., 2013). We focused on four sets of features: cortical thickness, cortical surface area, subcortical volume and total brain volume. For cortical thickness and cortical surface area, we used Destrieux’s atlas (Destrieux et al., 2010; Fischl, 2012) from FreeSurfer’s “aparc.stats” file, resulting in 148 regions for each set of features. For subcortical volume, we used the aseg atlas (Fischl et al., 2002) from FreeSurfer’s “aseg.stats” file, resulting in 19 regions. For total brain volume, we had five FreeSurfer-based features: “FS_IntraCranial_Vol” or estimated intra-cranial volume, “FS_TotCort_GM_Vol” or total cortical grey matter volume, “FS_Tot_WM_Vol” or total cortical white matter volume, “FS_SubCort_GM_Vol” or total subcortical grey matter volume and “FS_BrainSegVol_eTIV_Ratio” or ratio of brain segmentation volume to estimated total intracranial volume.

### Fluid cognition

We measured fluid cognition via the NIH Toolbox (Weintraub et al., 2014), using the “fluidcogcomp_unadj” variable. Fluid cognition summarises scores from five tests assessed outside of the MRI: Dimensional Change Card Sort, Flanker Inhibitory Control and Attention, Picture Sequence Memory, List Sorting Working Memory and Pattern Comparison Processing Speed.

### Prediction models for Brain Age and Brain Cognition

To compute Brain Age and Brain Cognition, we ran two separate prediction models. These prediction models either had chronological age or fluid cognition as the target and standardised brain MRI as the features (Denissen et al., 2022). We used nested cross-validation (CV) to build these prediction models (see Figure 8). We first split the data into five outer folds, leaving each outer fold with around 100 participants. This number of participants in each fold is to ensure the stability of the test performance across folds. In each outer-fold CV loop, one of the outer folds was treated as an outer-fold test set, and the rest was treated as an outer-fold training set. Ultimately, looping through the nested CV resulted in a) prediction models from each of the 18 sets of features as well as b) prediction models that drew information across different combinations of the 18 separate sets, known as “stacked models.” We specified eight stacked models: “All” (i.e., including all 18 sets of features), “All excluding Task FC”, “All excluding Task Contrast”, “Non-Task” (i.e., including only Rest FC and sMRI), “Resting and Task FC”, “Task Contrast and FC”, “Task Contrast” and “Task FC”. Accordingly, there were 26 prediction models in total for both Brain Age and Brain Cognition.

**Figure 8.**
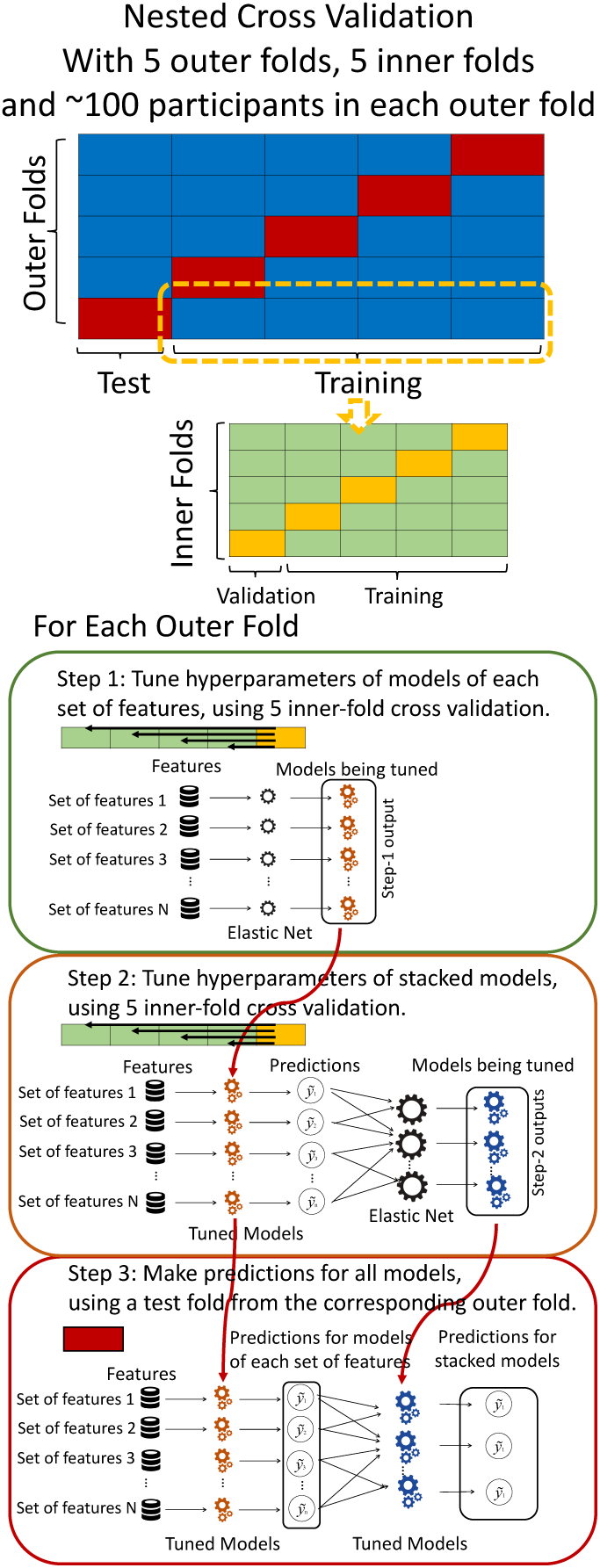
Diagram of the nested cross-validation used for creating predictions for models of each set of features as well as predictions for stacked models.

To create these 26 prediction models, we applied three steps for each outer-fold loop. The first step aimed at tuning prediction models for each of 18 sets of features. This step only involved the outer-fold training set and did not involve the outer-fold test set. Here, we divided the outer-fold training set into five inner folds and applied inner-fold CV to tune hyperparameters with grid search. Specifically, in each inner-fold CV, one of the inner folds was treated as an inner-fold validation set, and the rest was treated as an inner-fold training set. Within each inner-fold CV loop, we used the inner-fold training set to estimate parameters of the prediction model with a particular set of hyperparameters and applied the estimated model to the inner-fold validation set. After looping through the inner-fold CV, we, then, chose the prediction models that led to the highest performance, reflected by coefficient of determination (R^2^), on average across the inner-fold validation sets. This led to 18 tuned models, one for each of the 18 sets of features, for each outer fold.

The second step aimed at tuning stacked models. Same as the first step, the second step only involved the outer-fold training set and did not involve the outer-fold test set. Here, using the same outer-fold training set as the first step, we applied tuned models, created from the first step, one from each of the 18 sets of features, resulting in 18 predicted values for each participant. We, then, re-divided this outer-fold training set into new five inner folds. In each inner fold, we treated different combinations of the 18 predicted values from separate sets of features as features to predict the targets in separate “stacked” models. Same as the first step, in each inner-fold CV loop, we treated one out of five inner folds as an inner-fold validation set, and the rest as an inner-fold training set. Also as in the first step, we used the inner-fold training set to estimate parameters of the prediction model with a particular set of hyperparameters from our grid. We tuned the hyperparameters of stacked models using grid search by selecting the models with the highest R^2^ on average across the inner-fold validation sets. This led to eight tuned stacked models.

The third step aimed at testing the predictive performance of the 18 tuned prediction models from each of the set of features, built from the first step, and eight tuned stacked models, built from the second step. Unlike the first two steps, here we applied the already tuned models to the outer-fold test set. We started by applying the 18 tuned prediction models from each of the sets of features to each observation in the outer-fold test set, resulting in 18 predicted values. We then applied the tuned stacked models to these predicted values from separate sets of features, resulting in eight predicted values.

To demonstrate the predictive performance, we assessed the similarity between the observed values and the predicted values of each model across outer-fold test sets, using Pearson’s *r*, coefficient of determination (R^2^) and mean absolute error (MAE). Note that for R^2^, we used the sum of squares definition (i.e., R^2^ = 1 – (sum of squares residuals/total sum of squares)) per a previous recommendation (Poldrack et al., 2020). We considered the predicted values from the outer-fold test sets of models predicting age or fluid cognition, as Brain Age and Brain Cognition, respectively.

We controlled for the potential influences of biological sex on the brain features by first residualising biological sex from brain features in each outer-fold training set. We then applied the regression of this residualisation to the corresponding outer-fold test set. We also standardised the brain features in each outer-fold training set and then used the mean and standard deviation of this outer-fold training set to standardise the outer-fold test set. All of the standardisation was done prior to fitting the prediction models.

For the machine learning algorithm, we used Elastic Net (Zou & Hastie, 2005). Elastic Net is a general form of penalised regressions (including Lasso and Ridge regression), allowing us to simultaneously draw information across different brain indices to predict one target variable. Penalised regressions are commonly used for building age-prediction models (Jirsaraie, Gorelik, et al., 2023). Previously we showed that the performance of Elastic Net in predicting cognitive abilities is on par, if not better than, many non-linear and more-complicated algorithms (Pat, Wang, Bartonicek, et al., 2022; Tetereva et al., 2022). Moreover, Elastic Net coefficients are readily explainable, allowing us the ability to explain how our age-prediction and cognition-prediction models made the prediction from each brain feature (Molnar, 2019; Pat, Wang, Bartonicek, et al., 2022) (see below).

Elastic Net simultaneously minimises the weighted sum of the features’ coefficients. The degree of penalty to the sum of the feature’s coefficients is determined by a shrinkage hyperparameter ‘α’: the greater the α, the more the coefficients shrink, and the more regularised the model becomes. Elastic Net also includes another hyperparameter, ‘ℓ_1_ ratio’, which determines the degree to which the sum of either the squared (known as ‘Ridge’; ℓ_1_ ratio=0) or absolute (known as ‘Lasso’; ℓ_1_ratio=1) coefficients is penalised (Zou & Hastie, 2005). The objective function of Elastic Net as implemented by sklearn (Pedregosa et al., 2011) is defined as:

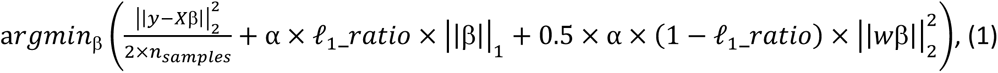

where *X* is the features, *y* is the target, and β is the coefficient. In our grid search, we tuned two Elastic Net hyperparameters: α using 70 numbers in log space, ranging from .1 and 100, and ℓ_1_-ratio using 25 numbers in linear space, ranging from 0 and 1.

To understand how Elastic Net made a prediction based on different brain features, we examined the coefficients of the tuned model. Elastic Net coefficients can be considered as feature importance, such that more positive Elastic Net coefficients lead to more positive predicted values and, similarly, more negative Elastic Net coefficients lead to more negative predicted values (Molnar, 2019; Pat, Wang, Bartonicek, et al., 2022). While the magnitude of Elastic Net coefficients is regularised (thus making it difficult for us to interpret the magnitude itself directly), we could still indicate that a brain feature with a higher magnitude weights relatively stronger in making a prediction. Another benefit of Elastic Net as a penalised regression is that the coefficients are less susceptible to collinearity among features as they have already been regularised (Dormann et al., 2013; Pat, Wang, Bartonicek, et al., 2022).

Given that we used five-fold nested cross validation, different outer folds may have different degrees of ‘α’ and ‘ℓ_1_ratio’, making the final coefficients from different folds to be different. For instance, for certain sets of features, penalisation may not play a big part (i.e., higher or lower ‘α’ leads to similar predictive performance), resulting in different ‘α’ for different folds. To remedy this in the visualisation of Elastic Net feature importance, we refitted the Elastic Net model to the full dataset without splitting them into five folds and visualised the coefficients on brain images using Brainspace (Vos De Wael et al., 2020) and Nilern (Abraham et al., 2014) packages. Note, unlike other sets of features, Task FC and Rest FC were modelled after data reduction via PCA. Thus, for Task FC and Rest FC, we, first, multiplied the absolute PCA scores (extracted from the ‘components_’ attribute of ‘sklearn.decomposition.PCA’) with Elastic Net coefficients and, then, summed the multiplied values across the 75 components, leaving 71,631 ROI-pair indices.

To demonstrate the stability of feature importance across outer folds, we examined the rank stability of feature importance using Spearman’s *ρ*. Specifically, we correlated the feature importance between two prediction models of the same features, used in two different outer-fold test sets. Given that there were five outer-fold test sets, we computed 10 Spearman’s *ρ* for each prediction model of the same features.

### Brain Age calculations: Brain Age, Brain Age Gap, Corrected Brain Age and Corrected Brain Age Gap

In addition to Brain Age, which is the predicted value from the models predicting chronological age in the outer-fold test sets, we calculated three other indices to reflect the estimation of brain aging. First, Brain Age Gap reflects the difference between the age predicted by brain MRI and the actual, chronological age. Here we simply subtracted the chronological age from Brain Age:

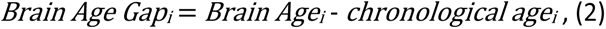

where i is the individual. Next, to reduce the dependency on chronological age (Butler et al., 2021; de Lange & Cole, 2020; Le et al., 2018), we applied a method described in de Lange and Cole’s (2020), which was implemented elsewhere (Cole et al., 2020; Cumplido-Mayoral et al., 2023; Denissen et al., 2022):

In each outer-fold training set:

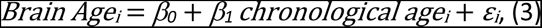

Then in the corresponding outer-fold test set:

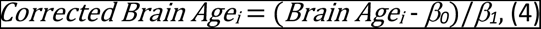

That is, we first fit a regression line predicting the Brain Age from a chronological age in each outer-fold training set. We then used the slope (β1) and intercept (β0) of this regression line to adjust Brain Age in the corresponding outer-fold test set, resulting in Corrected Brain Age. Note de Lange and Cole (2020) called this Corrected Brain Age, “Corrected Predicted Age”, while Butler (2021) called it “Revised Predicted Age.”

Lastly, we computed Corrected Brain Age Gap by subtracting the chronological age from the Corrected Brain Age (Butler et al., 2021; Cole et al., 2020; de Lange & Cole, 2020; Denissen et al., 2022):

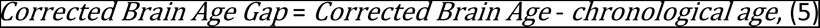

Note Cole and colleagues (2020) called Corrected Brain Age Gap, “brain-predicted age difference (brain-PAD),” while Butler and colleagues (2021) called this index, “Revised Brain Age Gap”.

### The utility of Brain Age indices to capture fluid cognition

We first combined Brain Age, Brain Cognition, chronological age and fluid cognition across outer-fold test sets into one table. We then conducted three sets of regression analyses to demonstrate the utility of different Brain Age indices, calculated from 26 different prediction models based on different sets of brain MRI features, to capture fluid cognition.

#### 1. Simple Regression: Using each Brain Age index to explain fluid cognition

Here using simple regression, we simply had each Brain Age index as the sole regressor for fluid cognition:

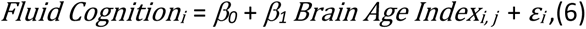

where *j* is the index for the four Brain Age indices. Because different Brain Age indices differ in the adjustments applied, this simple regression could reveal the extent to which each adjustment influences variation in fluid cognition explained. Additionally, Brain Age calculated from 26 different prediction models would have different levels of predictive performance in predicting chronological age. Accordingly, this simple regression could also reveal if Brain Age from a better-performing age-prediction model was able to capture more variation in fluid cognition.

In addition to Brain Age indices, we also used simple regression to test how well Brain Cognition as a sole regressor explains fluid cognition:

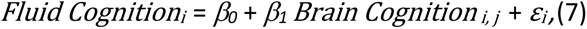

This allows us to compare the utility of Brain Age Indices vs. Brain Cognition as a sole regressor for predicting fluid cognition.

#### 2. Multiple Regression: Using chronological age and each Brain Age index to explain fluid cognition

Here using multiple regression, we had both chronological age and each Brain Age index as the regressors for fluid cognition:

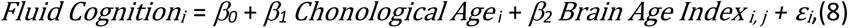

Having chronological age in the same regression model as a Brain Age index allowed us to control for the effects of chronological age on the Brain Age index, thereby, revealing the unique effects of the Brain Age index (Butler et al., 2021; Le et al., 2018). To formally determine the unique effects of a Brain Age index on fluid cognition along with the effects it shared with chronological age (i.e., common effects), we applied the commonality analysis (Nimon et al., 2008). For the unique effects, we computed ΔR^2^. ΔR^2^ is the increase in R^2^ when having an additional regressor in the regression model:

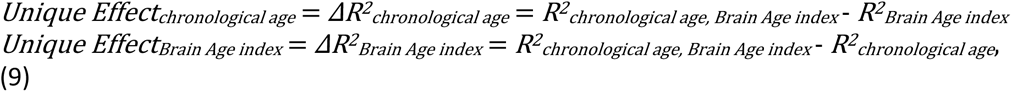

We determined the statistical significance of ΔR^2^ by:

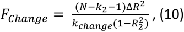

where *F_Change_*is the *F*-ratio (with the degree of freedom of *k_Change_* and *N* – *k_2_* –1), *N* is the number of observations, _2_ is the model with more regressors, *k* is the number of regressors, *k_Change_* is the difference between the number of regressors.

As for the common effects between chronological age and each Brain Age index, we used the below calculation:

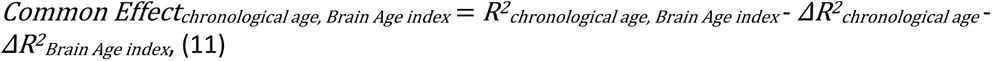

These common effects indicate the extent to which variation in fluid cognition explained by each Brain Age index was shared with chronological age. Note the common effects can be negative, especially with a high multicollinearity (Ray-Mukherjee et al., 2014). To deal with this, we treated negative common effects as zero (Frederick, 1999) and then scaled variation explained by other effects to be proportional to the total effects of the full regression model.

#### 3. Multiple Regression: Using chronological age, each Brain Age index and Brain Cognition to explain fluid cognition

Similar to the above multiple regression model, we had chronological age, each Brain Age index and Brain Cognition as the regressors for fluid cognition:

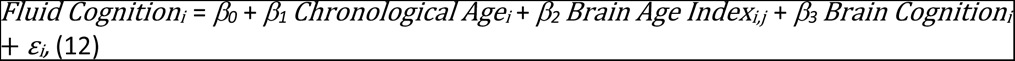

Applying the commonality analysis here allowed us, first, to investigate the addictive, unique effects of Brain Cognition, over and above chronological age and Brain Age indices. More importantly, the commonality analysis also enabled us to test the common, shared effects that Brain Cognition had with chronological age and Brain Age indices in explaining fluid cognition. We calculated the commonality analysis as follows (Nimon et al., 2017):

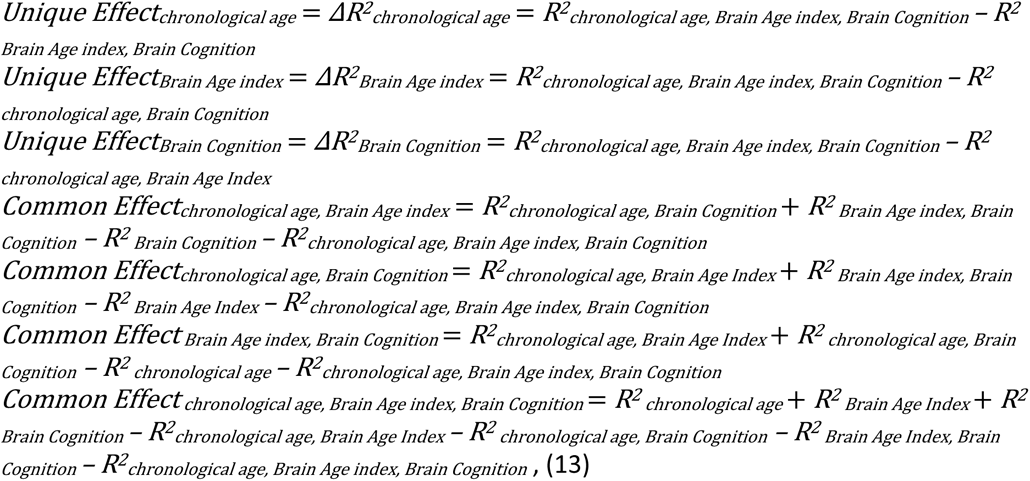

Note to ensure that the commonality analysis results were robust against multicollinearity (Ray-Mukherjee et al., 2014), we also repeated the same commonality analyses done here on Ridge regression, as opposed to multiple regression. Ridge regression is a method designed to deal with multicollinearity (Dormann et al., 2013). See Supplementary Figure 3 for the Ridge regression with chronological age and each Brain Age index as regressors and Supplementary Figure 5 for the Ridge regression with chronological age, each Brain Age and Brain Cognition index as regressors. Briefly, the results from commonality analyses applied to Ridge regressions are closely matched with our results done using multiple regression.

Similarly, to ensure that we were able to capture the non-linear pattern of chronological age in explaining fluid cognition, we added a quadratic term of chronological age to our multiple-regression models in the commonality analyses. See Supplementary Figure 4 for the multiple regression with chronological age, square chronological age and each Brain Age index as regressors and Supplementary Figure 6 for the multiple regression with chronological age, square chronological age, each Brain Age index and Brain Cognition as regressors. Briefly, adding the quadratic term for chronological age did not change the pattern of the results of the commonality analyses.

## Acknowledgements

Data were provided by the Human Connectome Project in Aging. Research reported in this publication was supported by the National Institute On Aging of the National Institutes of Health under Award Number U01AG052564 and by funds provided by the McDonnell Center for Systems Neuroscience at Washington University in St. Louis. The content is solely the responsibility of the authors and does not necessarily represent the official views of the National Institutes of Health. The author(s) wish to acknowledge the use of New Zealand eScience Infrastructure (NeSI) high performance computing facilities, consulting support and/or training services as part of this research. New Zealand’s national facilities are provided by NeSI and funded jointly by NeSI’s collaborator institutions and through the Ministry of Business, Innovation & Employment’s Research Infrastructure programme. URL https://www.nesi.org.nz. A.T. and N.P. were supported by Health Research Council Funding (21/618) and by the University of Otago.

## Code Accessibility

The shell and Python scripts used in the analyses are made available here: https://github.com/HAM-lab-Otago-University/HCP-Aging_commonality

## Supplementary

**Supplementary Figure 1.**
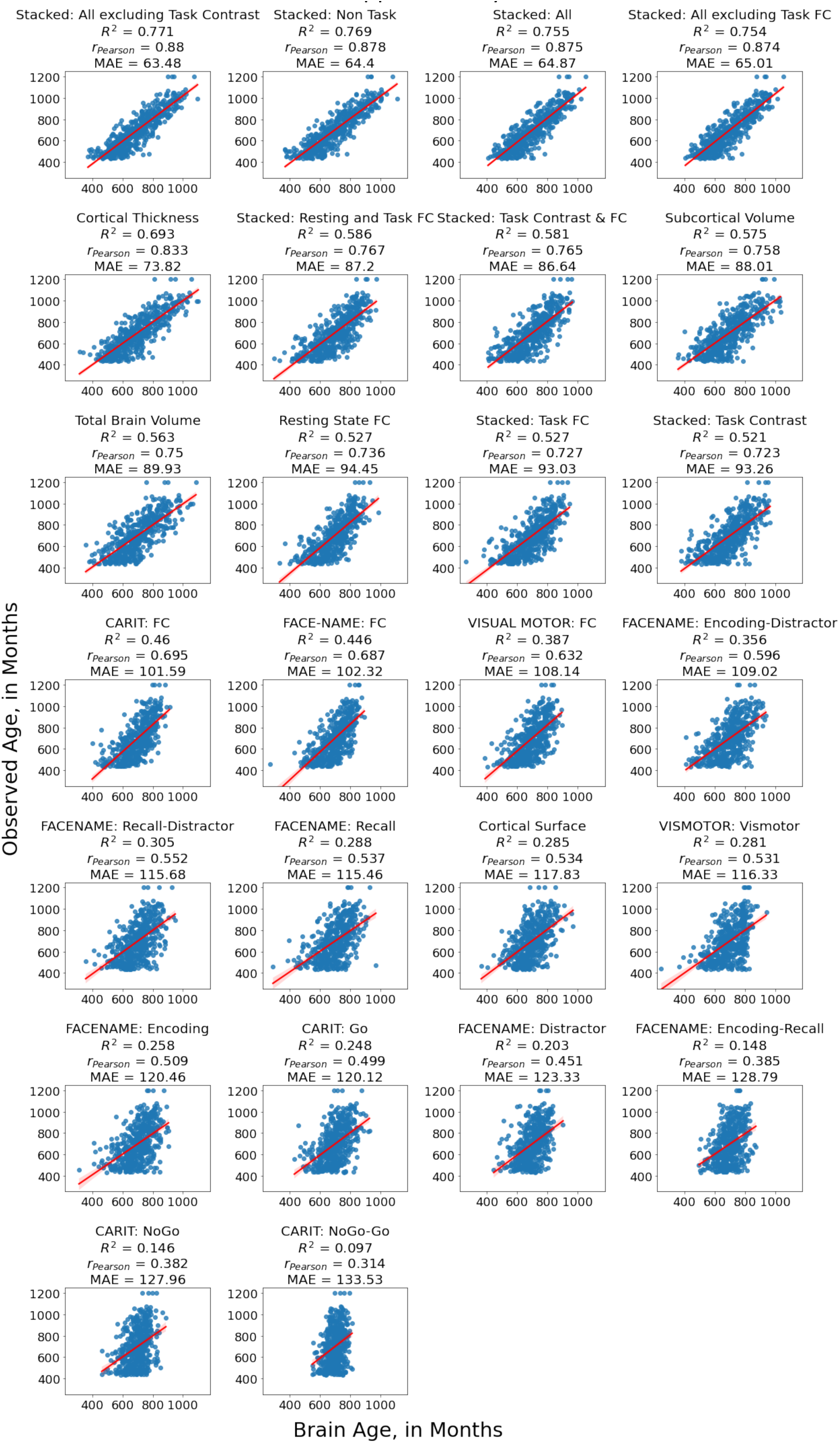
The scatter plots between observed and predicted values in the outer-fold test sets from age-prediction models.

**Supplementary Figure 2.**
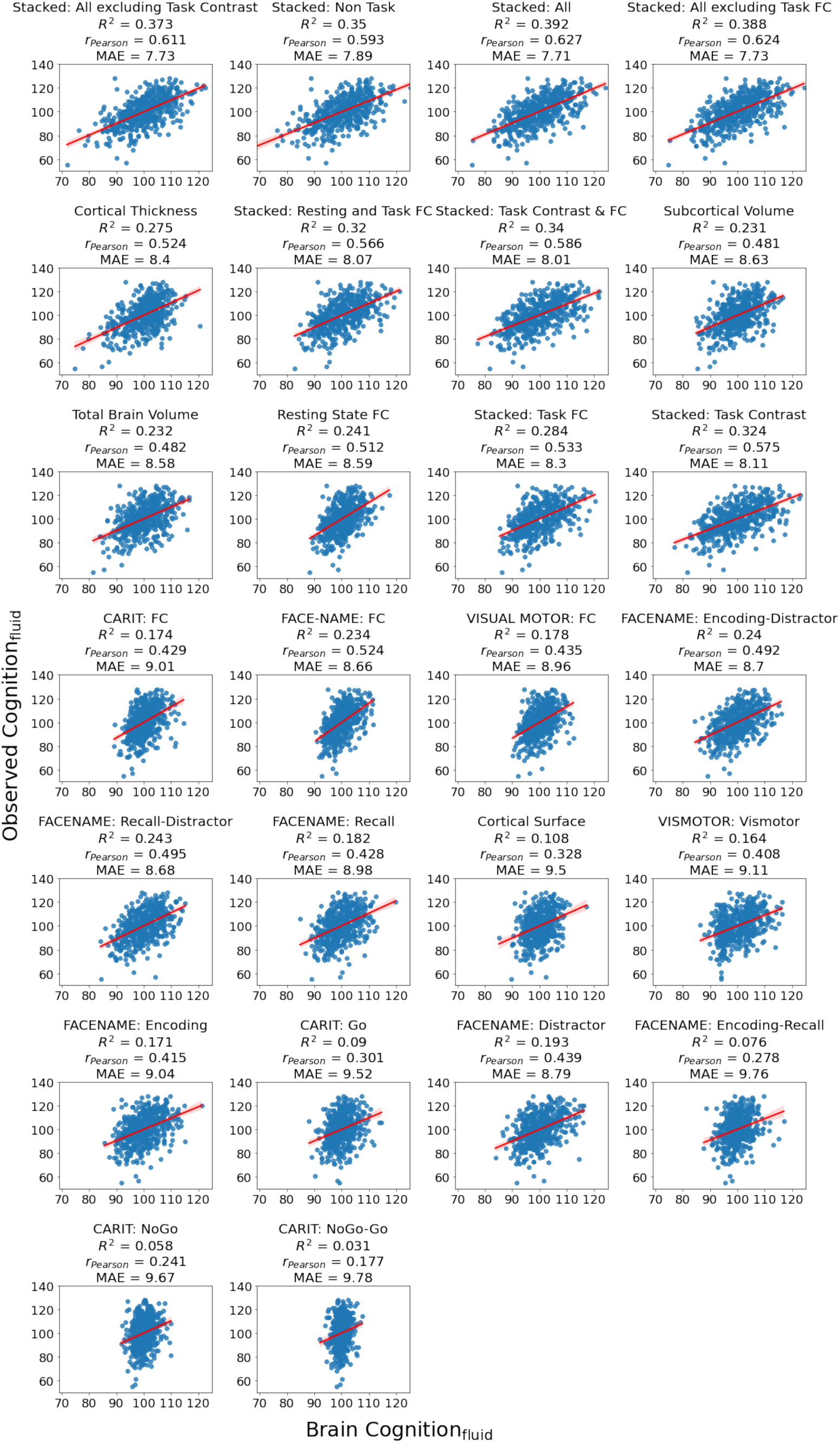
The scatter plots between observed and predicted values in the outer-fold test sets from cognition-prediction models.

**Supplementary Figure 3.**
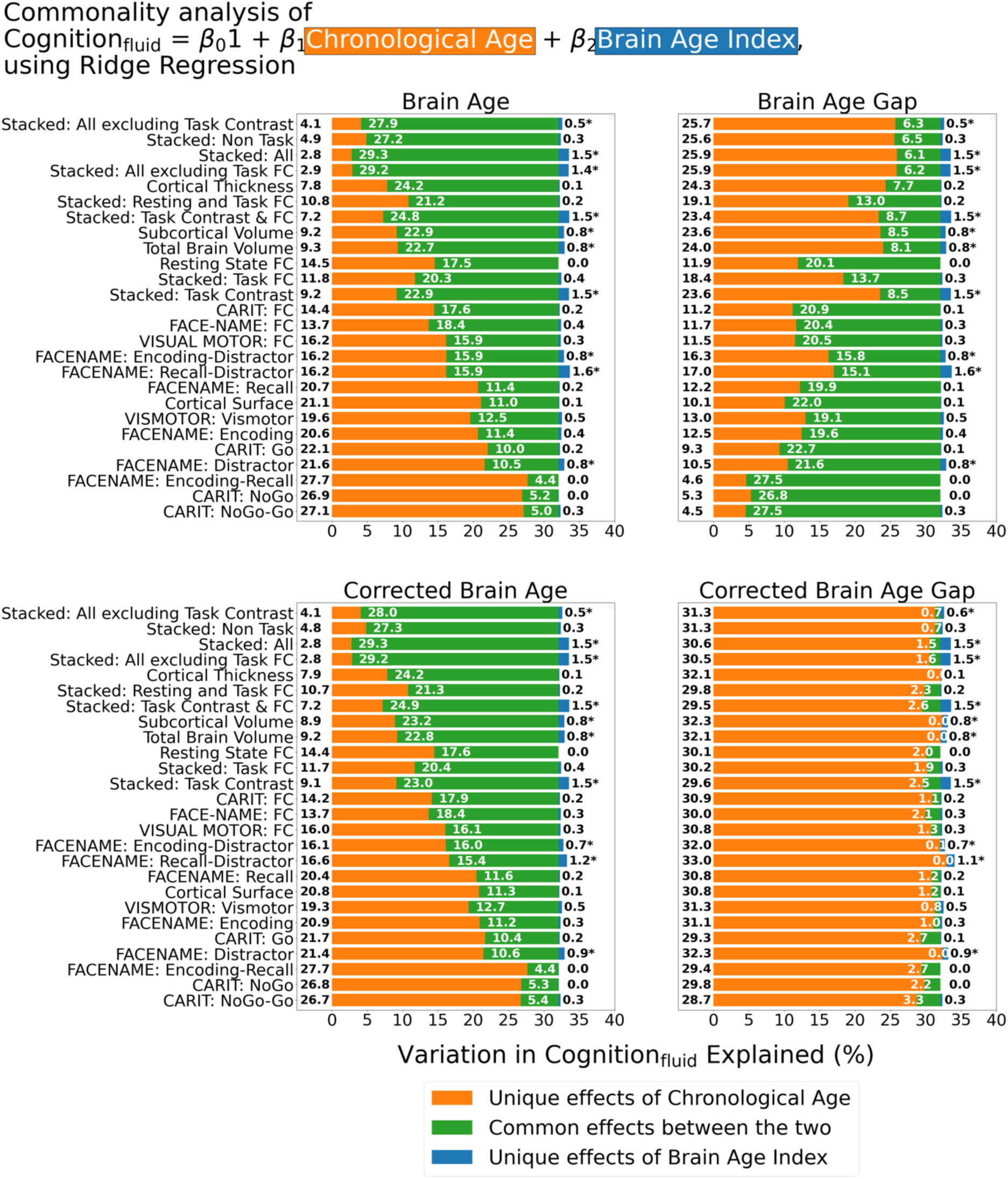
Commonality analysis of Ridge regressions, having both chronological age and each Brain Age index as the regressors for capturing fluid cognition. Note we used Ridge regressions for models with both chronological age and each Brain Age index as the regressors and simple regressions for models with a single regressor (apart from the intercept). The numbers to the left of the figures represent the unique effects of chronological age in %, the numbers in the middle of the figures represent the common effects between chronological age and Brain Age index in %, and the numbers to the right of the figures represent the unique effects of Brain Age Index in %. * represents the statistical significance of the unique effects of Brain Age Index at p < .05.

**Supplementary Figure 4.**
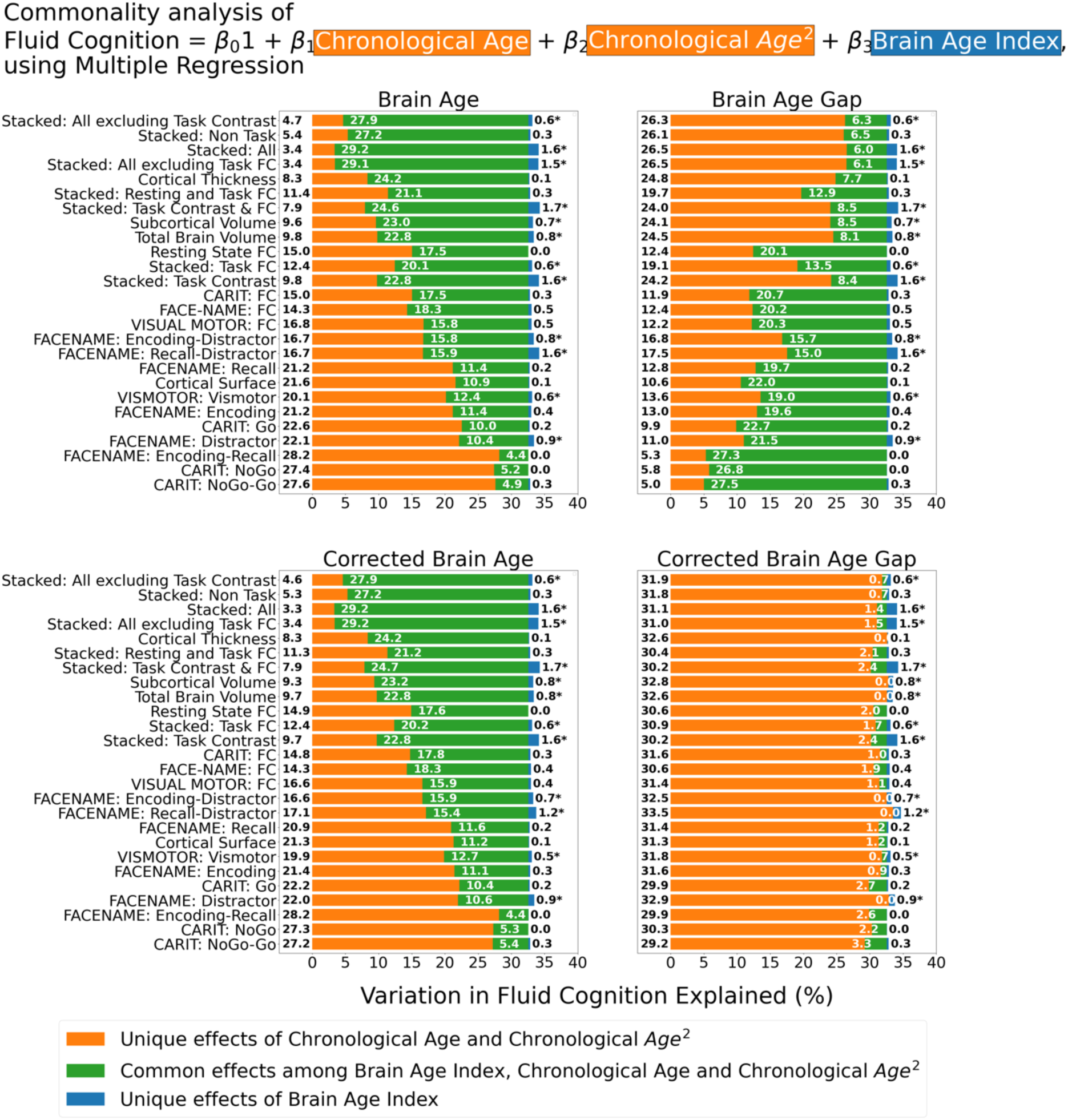
Commonality analysis of multiple regressions, having chronological age, a quadratic term for chronological age and each Brain Age index as the regressors for fluid cognition. The numbers to the left of the figures represent the unique effects of chronological age in %, the numbers in the middle of the figures represent the common effects between chronological age and Brain Age index in %, and the numbers to the right of the figures represent the unique effects of Brain Age Index in %. * represents the statistical significance of the unique effects of Brain Age Index at p < .05.

**Supplementary Figure 5.**
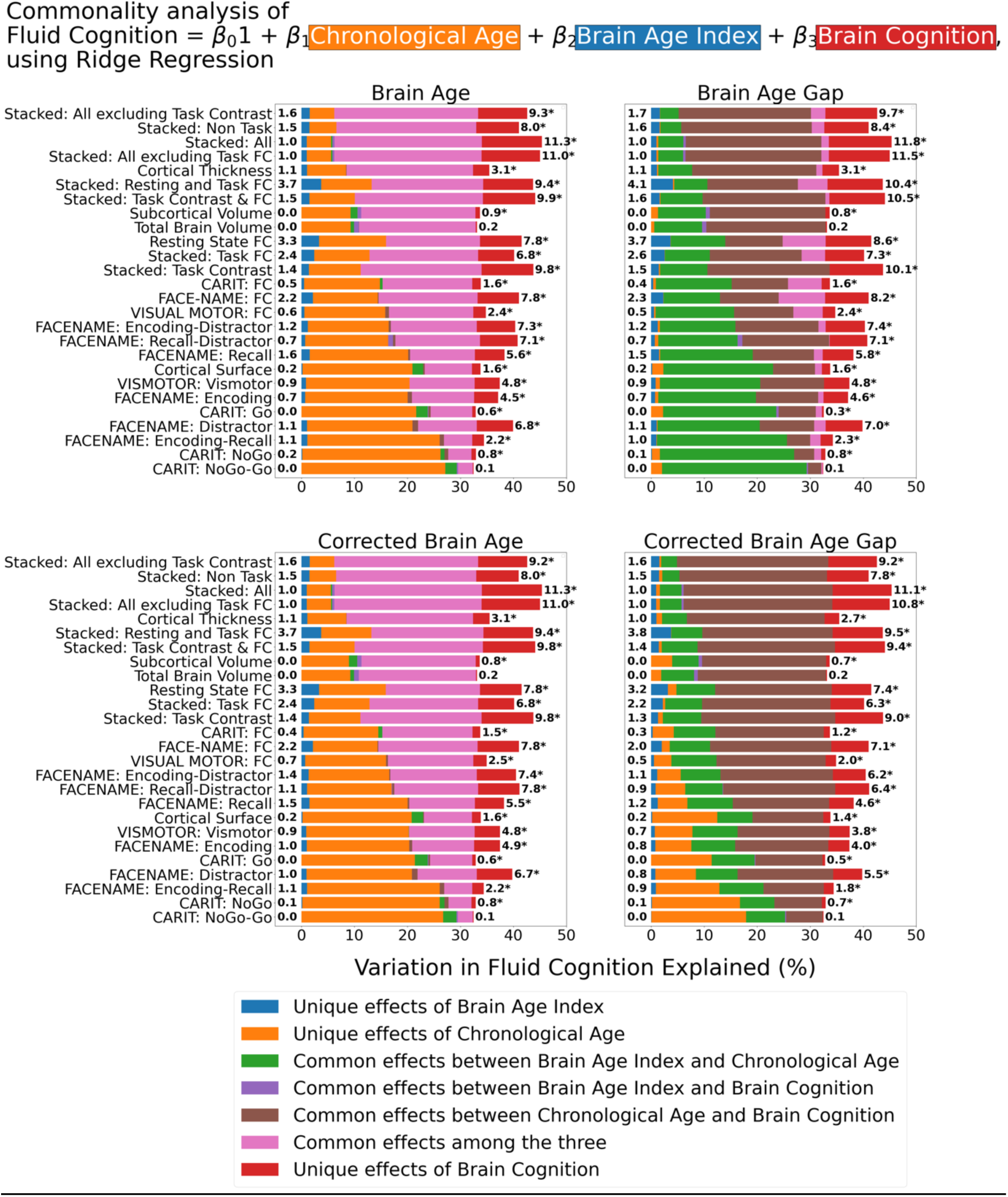
Commonality analysis of Ridge regressions, having chronological age and each Brain Age index and Brain Cognition as the regressors for fluid cognition. Note we used Ridge regressions for models with at least two regressors (apart from the intercept) and simple regressions for models with a single regressor (apart from the intercept). The numbers to the left of the figures represent the unique effects of Brain Age Index in %, and the numbers to the right of the figures represent the unique effects of Brain Cognition in %. * represents the statistical significance of the unique effects of Brain Cognition at p < .05.

**Supplementary Figure 6.**
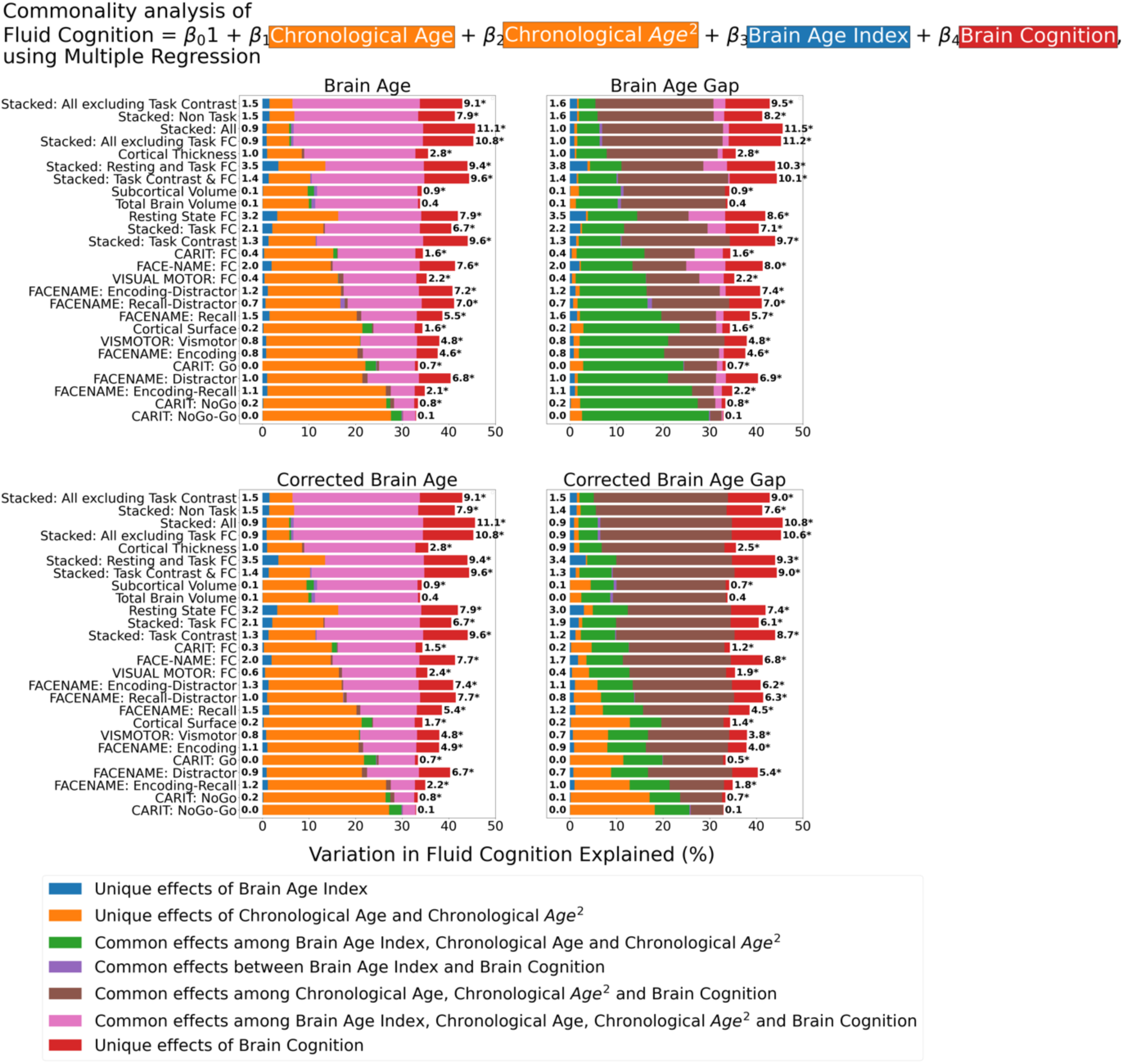
Commonality analysis of multiple regressions, having chronological age, a quadratic term for chronological age and each Brain Age index and Brain Cognition as the regressors for fluid cognition. The numbers to the left of the figures represent the unique effects of Brain Age Index in %, and the numbers to the right of the figures represent the unique effects of Brain Cognition in %. * represents the statistical significance of the unique effects of Brain Cognition at p < .05.

